# Spike burst–pause dynamics of Purkinje cells regulate sensorimotor adaptation

**DOI:** 10.1101/347252

**Authors:** Niceto R. Luque, Francisco Naveros, Richard R. Carrillo, Eduardo Ros, Angelo Arleo

## Abstract

Cerebellar Purkinje cells mediate accurate eye movement coordination. However, it remains unclear how oculomotor adaptation depends on the interplay between the characteristic Purkinje cell response patterns, namely tonic, bursting, and spike pauses. Here, a spiking cerebellar model assesses the role of Purkinje cell firing patterns in vestibular ocular reflex (VOR) adaptation. The model captures the cerebellar microcircuit properties and it incorporates spike-based synaptic plasticity at multiple cerebellar sites. A detailed Purkinje cell model reproduces the three spike-firing patterns that are shown to regulate the cerebellar output. Our results suggest that pauses following Purkinje complex spikes (bursts) encode transient disinhibition of targeted medial vestibular nuclei, critically gating the vestibular signals conveyed by mossy fibres. This gating mechanism accounts for early and coarse VOR acquisition, prior to the late reflex consolidation. In addition, properly timed and sized Purkinje cell bursts allow the ratio between long-term depression and potentiation (LTD/LTP) to be finely shaped at mossy fibre-medial vestibular nuclei synapses, which optimises VOR consolidation. Tonic Purkinje cell firing maintains the consolidated VOR through time. Importantly, pauses are crucial to facilitate VOR phase-reversal learning, by reshaping previously learnt synaptic weight distributions. Altogether, these results predict that Purkinje spike burst-pause dynamics are instrumental to VOR learning and reversal adaptation.

**Author Summary:** Cerebellar Purkinje cells regulate accurate eye movement coordination. However, it remains unclear how cerebellar-dependent oculomotor adaptation depends on the interplay between Purkinje cell characteristic response patterns: tonic, high-frequency bursting, and post-complex spike pauses. We explore the role of Purkinje spike burst-pause dynamics in VOR adaptation. A biophysical model of Purkinje cell is at the core of a spiking network model, which captures the cerebellar microcircuit properties and incorporates spike-based synaptic plasticity mechanisms at different cerebellar sites. We show that Purkinje spike burst-pause dynamics are critical for (1) gating the vestibular-motor response association during VOR acquisition; (2) mediating the LTD/LTP balance for VOR consolidation; (3) reshaping synaptic efficacy distributions for VOR phase-reversal adaptation; (4) explaining the reversal VOR gain discontinuities during sleeping.

## Introduction

The cerebellum controls fine motor coordination including online adjustments of eye movements [1]. Within the cerebellar cortex, the inhibitory projections of Purkinje cells to medial vestibular nuclei (MVN) mediate the acquisition of accurate oculomotor control [2, 3]. Here, we consider the role of cerebellar Purkinje cells in the adaptation of the vestibular ocular reflex (VOR), which generates rapid contralateral eye movements that maintain images in the fovea during head rotations (Fig 1A). The VOR is crucial to preserve clear vision (e.g., whilst reading) and maintain balance by stabilising gaze during head movements. The VOR is mediated by the three-neuron reflex arc comprised of connections from the vestibular organ via the medial vestibular nuclei (MVN) to the eye motor neurons [3–5]. VOR control is purely feed-forward [6] and it relies on several cerebellar-dependent adaptive mechanisms driven by sensory errors (Fig 1B). Because of its dependence upon cerebellar adaptation, VOR has become one of the most intensively used paradigms to assess cerebellar learning [6]. However, very few studies have focused on the relation between the characteristics spike response patterns of Purkinje cells and VOR adaptation, which is the main focus of this study.

**Figure 1.**
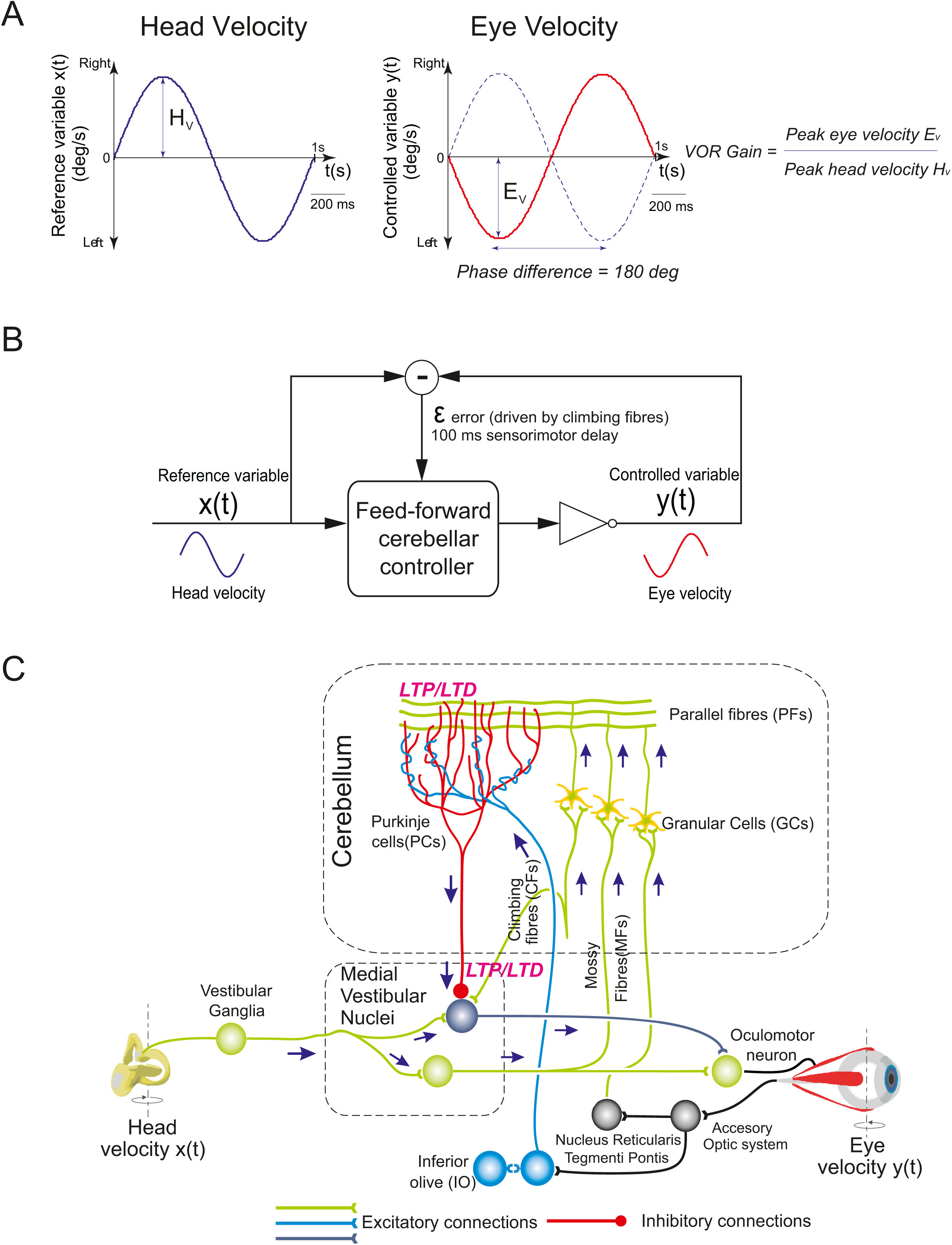
Vestibular Ocular Reflex (VOR) and cerebellar control loop. **(A)** Horizontal VOR (h-VOR) protocols compare head rotational movements (input) against the induced contralateral eye movements (output) via two measurements: the VOR gain, i.e. the ratio between eye and head speeds (E_v_ and H_v_, respectively); and the VOR phase, i.e. the temporal lag between eye and head velocity signals. **(B)** Cerebellar feed-forward control system comparing a known reference (head velocity or input variable) to the actual output (eye velocity) to quantify an error signal driving adaptation. The cerebellum compensates for the difference between actual eye (represented as an inverter logic gate in this scheme) and head velocity profiles. The head velocity consists of a 1 Hz sinusoidal function iteratively presented to the cerebellar model, mimicking the sinusoidal frequency of the head rotation in experimental protocols [7]. **(C)** Schematic representation of the main neural layers, cells, connections and plasticity sites considered in the cerebellar model. Mossy fibres (MFs) convey the sensory signals from the vestibular organ and they provide the input to the cerebellar network. MFs project sensorimotor information onto granular cells (GCs) and medial vestibular nuclei (MVN). GCs, in turn, project onto Purkinje cells through parallel fibres (PFs). Purkinje cells also receive excitatory inputs from the climbing fibres (CFs). CFs deliver the error signals encoding instructive terms that drive motor control learning. Purkinje cells integrate CF and PF inputs, thus transmitting the difference between head and eye movements. Finally, MVN are inhibited by Purkinje cells and provide the main cerebellar output. The cerebellar model implements different spike timing dependent plasticity mechanisms at multiple sites: PF-Purkinje cell, MF-MVN, and Purkinje cell-MVN synapses.

Purkinje cells provide the major output of the cerebellum through MVN. Purkinje cells receive two main excitatory (glutamatergic) afferent currents (Fig 1C). The first excitatory input originates from the parallel fibres (PFs), i.e. the axons of the granule cells (GCs). The second comes from the climbing fibres (CFs), i.e. the projections of the inferior olive (IO) cells. These excitatory inputs drive Purkinje cell simple or complex spike patterns, respectively [8, 9]. Simple spikes of Purkinje cells are elicited topically at high frequencies [10, 11]. Complex spikes consist of a fast initial large-amplitude spike followed by a high-frequency burst [12]. This burst is made of several slower spikelets of smaller amplitude separated from one another by 2-3 ms [12–14]. Complex spikes are caused by the activation of a single IO neuron that produces a large electrical event in the soma of the post-synaptic Purkinje cell. This electrical event generates calcium-mediated action potentials in the Purkinje cell dendrites that, in turn, shape the complex spike. Simple spike activity is, in fact, mostly suppressed during complex spiking [14]. After each CF-evoked burst, a spike pause prevents Purkinje cells from either resuming their tonic or bursting firing for a period that depends on the length of the complex spike [15]. The CF-evoked spike burst-pause sequences of Purkinje cell responses critically regulate the inhibitory (GABAergic) drive of MVN synapses, which determines the cerebellar output during sensorimotor adaptation. Therefore, understanding the dynamics of the characteristic Purkinje cell spike patterns is relevant to linking cerebellar cell properties to cerebellar-dependent behavioural adaptation. Recent studies have paved the road in gaining knowledge on the behavioural implication of Purkinje cell spike modes [2, 14, 16]. In particular, Herzfeld and colleagues have demonstrated that the cerebellum encodes real-time motion of the eye through the organisation of Purkinje cells into clusters that share similar CF projections from the IO [2]. The combined activity of bursting and silent Purkinje cell populations can predict both the actual speed and direction of rapid accurate eye movements (saccades). However, these studies have not assessed the interplay between the different Purkinje cell spike patterns and the plasticity mechanisms at stake at MVN synapses in shaping sensorimotor adaptation. MVN neurons, in addition to receiving the inhibitory inputs from Purkinje cells, are also innervated by the excitatory afferents from the mossy fibres (MFs), which convey vestibular signals about head movements (Fig 1C). This vestibular information also converges onto Purkinje cells through the mossy fibre-granule cell-parallel fibre pathway (MF-GC-PF; Fig 1C). Therefore, the characteristics firing patterns of Purkinje cells are likely to play a key role in driving the associative plasticity mechanisms operating at MF-MVN excitatory synapses [17–19] and at Purkinje cells-MVN inhibitory synapses [20–23]. The CF-evoked spike burst-pause sequences of Purkinje cells depend indeed upon the activation of CFs, which are assumed to convey a ‘teaching’ signal encoding sensory error information [6, 14, 24]. Therefore, the properties of the CF-evoked spike burst-pause patterns (e.g., the relative duration of the bursts versus the pauses) reflect sensory error related information [14, 16]. The activation of CFs is critical for inducing different forms of plasticity at PF-Purkinje cell synapses and, indirectly, at Purkinje cell-MVN synapses [25, 26]. Importantly, plasticity at MF-MVN synapses also seems to be dependent on Purkinje cell signals [27–29], generated through the MF-GC-PF pathway and through CF activation. Some computational studies have proposed that plasticity mechanisms at MF-MVN and Purkinje cell-MVN synapses are key factors in determining cerebellar adaptive gain control [27, 28, 30]. These models support the hypothesis of a two-state cerebellar adaptation process [31, 32], with a fast adaptive phase mediated by the cerebellar cortex (involving plasticity at Purkinje cell synapses) and a slow adaptive process occurring in deeper structures, involving plasticity at MVN synapses [29, 31–35]. However, these computational studies do not account for the interaction between the different spiking modes of Purkinje cells (in particular CF-evoked spike burst-pause dynamics) and the distributed plasticity mechanisms underpinning cerebellar adaptive control [30].

The spiking cerebellar model presented here addresses these issues within a VOR adaptation framework (Figs 1A,B). We simulate horizontal VOR (h-VOR) experiments with mice undertaking sinusoidal (∼1 Hz) whole body rotations in the dark [36]. The model incorporates the main anatomo-functional properties of the cerebellar microcircuit, with synaptic plasticity mechanisms at multiple cerebellar sites (Fig 1C; see Materials & Methods).

## Results

### Spike burst–pause properties of model Purkinje cell responses

The detailed Purkinje cell model reproduces the characteristic response patterns observed experimentally: tonic simple spiking (20-200 Hz), complex spiking (bursts with high-frequency spikelet components up to 600 Hz), and post-complex spike pauses (Fig 2A). In the model, CF discharges trigger transitions between the Purkinje cell Na^+^ spike output, CF-evoked bursts, and post-complex spike pauses. As evidenced in [37], in *in-vitro* slice preparations at normal physiological conditions, 70% of Purkinje cells spontaneously express a trimodal oscillation: a Na^+^ tonic spike phase, a Ca-Na^+^ bursting phase, and a hyperpolarised quiescent phase. On the other hand, Purkinje cells also show spontaneous firing consisting of a tonic Na^+^ spike output without Ca-Na^+^ bursts [37–39]. McKay et al. [37] report Purkinje cell recordings exhibiting a tonic Na^+^ phase sequence followed by CF-evoked bursts (via complex spikes) and the subsequent pause (Fig 2A). The frequency of Purkinje cell Na^+^ spike output decreases with no correlation with the intervals between CF discharges. The model mimics this behaviour under similar CF discharge conditions (Fig 2B).

**Figure 2.**
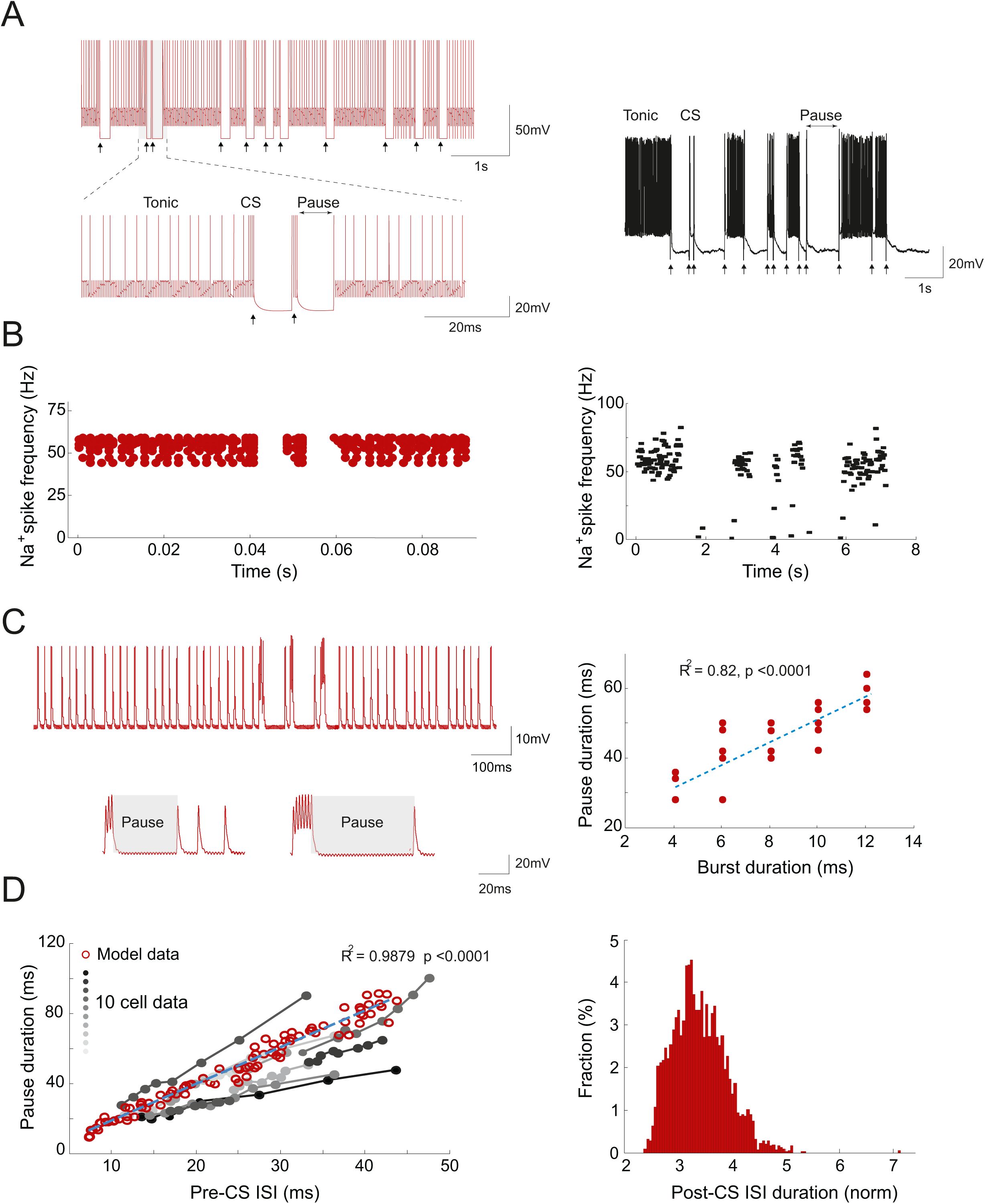
Spike burst–pause properties of model Purkinje cell responses. **(A)** Simulated (left) and electrophysiological (right) recordings of Purkinje cell spike outputs in response to CF spike excitatory postsynaptic potentials occurring at physiological frequencies (arrows) (data from [37]). CF discharges trigger transitions between Purkinje cell Na^+^ spike output and CF-evoked bursts and pauses via complex spikes. Here, the Purkinje cell model was run on the EDLUT simulator (see Methods). **(B)** Simulated (left) and experimental (right) Purkinje cell tonic spike frequency during CF discharges aligned with spike-grams in A (data from [37]). N=10 Purkinje cells were simulated to compute the tonic spike frequency. **(C)** In the model, CF signals modulate both the burst size (i.e., the number of spikes within the burst) and the duration of post-complex spike pauses, which are linearly correlated. Here, the Purkinje cell model was run on the Neuron simulator (see Methods). **(D)** Relation between pause duration and pre-complex spike (pre–CS) inter spike intervals (ISIs) when increasing the amplitude of the injected current: model data (red circles, n=1000) vs. experimental data [40] (grey to black dots). Grey-to-black lines represent individual cells (n=10). The blue dashed line is the linear regression curve fitting model data. The model captures the linear relation between spike pause duration and pre-complex spike ISI duration observed electrophysiologically [40]. **(E)** Distribution of ISI values following the complex spike (post-CS). The ISI duration is normalised to pre-CS ISI values. The Kurtosis for the distribution of post-CS ISI values is 4.24. The skewness is positive (0.6463), thus indicating an asymmetric post-CS ISI distribution. Kurtosis and skewness values were consistent with Purkinje cell data [40].

The duration of model post-complex spike pauses increases linearly with burst duration (Fig 2C; R^2^=0.82, p<0.0001). To assess the relation between burst and pause duration, the depolarisation current injected through PF was maintained constant whilst progressively increasing the intensity of CF stimulation. Only inter-spike intervals (ISIs) immediately following complex spikes were considered for this analysis. The model replicates the linear relation between spike pause duration and *pre-*complex spike ISI duration observed through electrophysiological recordings [40] (Fig 2D; R^2^=0.9879; p<0.0001). This relation was measured by maintaining the CF stimulation constant whilst incrementally increasing the amplitude of the PF input current. The probability distribution of *post*-complex spike ISIs is also consistent with experimental data [40] (Fig 2E). The kurtosis (‘peakedness’) of the ISI distribution is 4.24, which is in the range of kurtosis values measured after tetanisation of mouse Purkinje cells [40]. Finally, model *post*-complex spike ISI values are skewed rightward (positive skewness value of 0.6463), consistently with the asymmetric distribution shape observed experimentally [40].

### Role of cerebellar Purkinje spike burst-pause dynamics in VOR adaptation

We assessed h-VOR adaptation by simulating a 1 Hz horizontal head rotation to be compensated by contralateral eye movements (Fig 1A). First, we tested the role of Purkinje spike burst-pause dynamics in the absence of cerebellar learning, i.e. by blocking synaptic plasticity across all model projections (i.e., MF-MVN, PF-Purkinje cell, Purkinje cell-MVN). Synaptic weights were initialised randomly and equally within each projection set. The CF input driving Purkinje cells was taken as to signal large retina slips, which generated sequences of complex spikes made of 4 to 6 burst spikelets [14] (Fig 3A, top). The elicited Purkinje spike burst-pause sequences shaped the temporal disinhibition of targeted VN neurons, allowing the incoming input from MFs to drive MVN responses (Fig 3A, middle). This facilitated a coarse baseline eye motion (Fig 3A, bottom). Blocking complex spiking in the Purkinje cell model (through the blockade of muscarinic voltage-dependent channels, see Methods) prevented MF activity from eliciting any baseline MVN compensatory output (Fig 3B). These results suggest that the gating mechanism mediated by Purkinje spike burst-pause sequences, which encode transient disinhibition of MVN neurons, is useful for early and coarse VOR, prior to the adaptive consolidation of the reflex through cerebellar learning.

**Figure 3.**
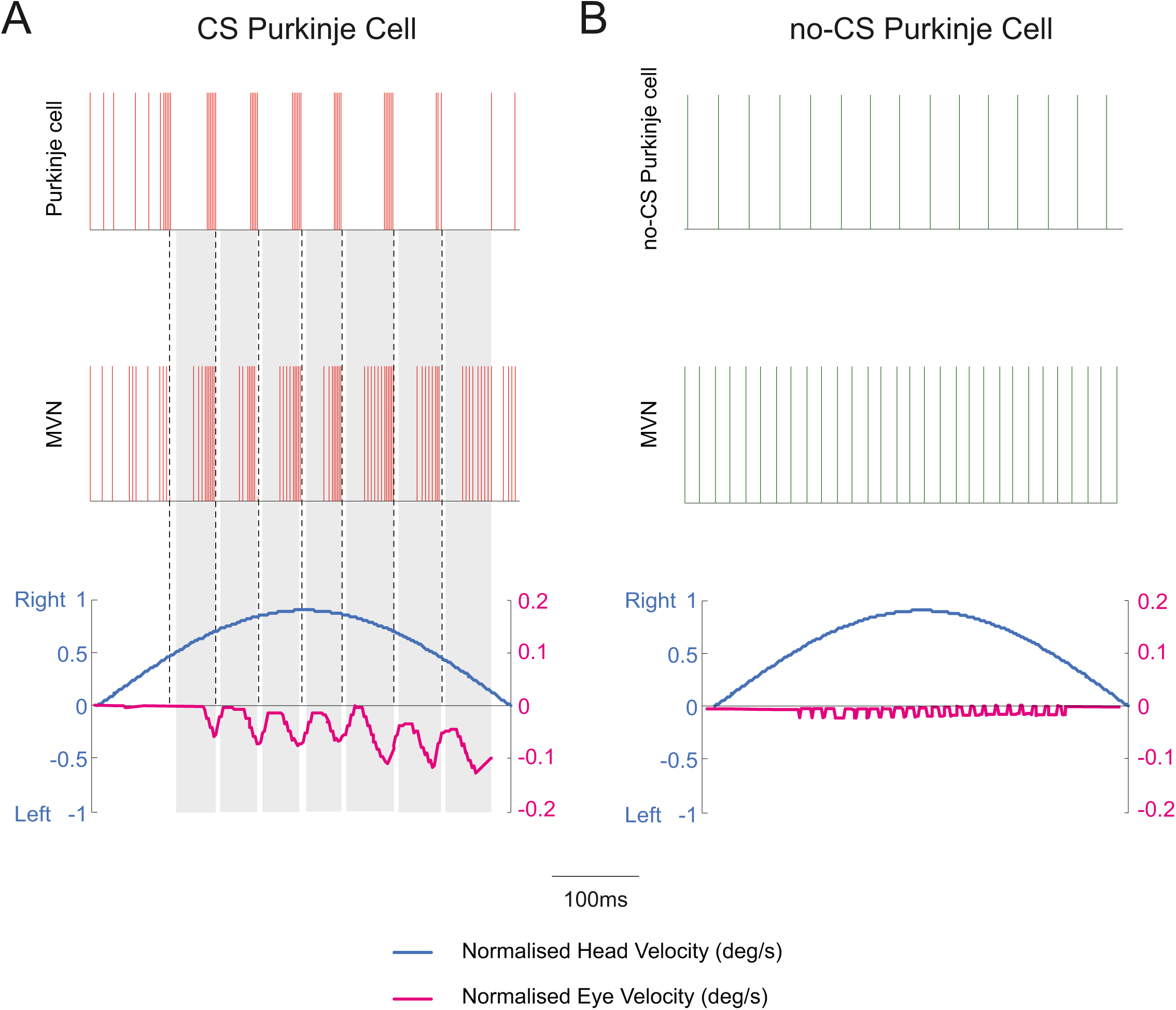
Purkinje post–complex spike pauses act as a gating mechanism for early coarse VOR in the absence of cerebellar adaptation. Only half of h-VOR cycle is represented. Two equal cerebellar network configurations except for the Purkinje cell dynamics were compared under equal stimulation. **(A)** The first model accounts for CF-evoked Purkinje spike burst-pause dynamics. CF stimulation generates complex spikes and subsequent post–complex spike pauses. The latter allows MFs to drive directly the immediate activation of MVN, which facilitates an early but rough eye movement compensation for head velocity. **(B)** The second model only exhibits Purkinje tonic firing (i.e., complex spiking is blocked through the blockade of muscarinic voltage-dependent channels, see Methods), which prevents MFs from eliciting any baseline MVN compensatory output. See S3-1 and S3-2 Figs for a sensitivity analysis of parameters regulating the LTD/LTP balance at PF-Purkinje cell and MF-MVN synapses. See also S3-3 Fig for the same parameter sensitivity analysis in the absence of Purkinje spike burst-pause dynamics.

We then activated the LTD/LTP plasticity mechanisms at MF-MVN, PF-Purkinje cell, and Purkinje cell-MVN synapses (see Materials & Methods). During 10000 s, the model faced a 1 Hz horizontal head rotation, and cerebellar h-VOR learning took place to generate compensatory contralateral eye movements. A sensitivity analysis identified the critical LTD/LTP balance at MF-MVN and PF-Purkinje cell synapses in order to achieve VOR adaptation (in terms of both gain and phase). This analysis predicts a very narrow range of values for which LTP slightly exceeding LTD at MF-MVN synapses ensures learning stability through time. By contrast, PF-Purkinje cell synapses admitted a significantly broader range for the LTD/LTP ratio (S3-1 and S3-2 Figs). The same parameter sensitivity analysis for the cerebellar model with no bursting and pause dynamics shows a much wider range of values for the LTD/LTP balance at both PF-Purkinje cell and MF-MVN synapses (S3-3 Fig).

A comparison of VOR adaptation accuracy in the presence vs. absence of CF-evoked Purkinje spike burst-pause dynamics shows that VOR gain plateaued three times faster in the presence of Purkinje complex spikes (Fig 4A, left). Also, the VOR gain converged to [0.8-0.9], which is consistent with experimental recordings in mice [36], monkeys [41], and humans [42]. Conversely, without Purkinje bursting-pause dynamics the VOR gain saturated to a value >1 (i.e., over learning) at the end of the adaptation process. In terms of VOR phase, convergence to 180° (i.e., well synchronised counter-phase eye movements) was reached after approximately 1000 s under both conditions (Fig 4A, right).

**Figure 4.**
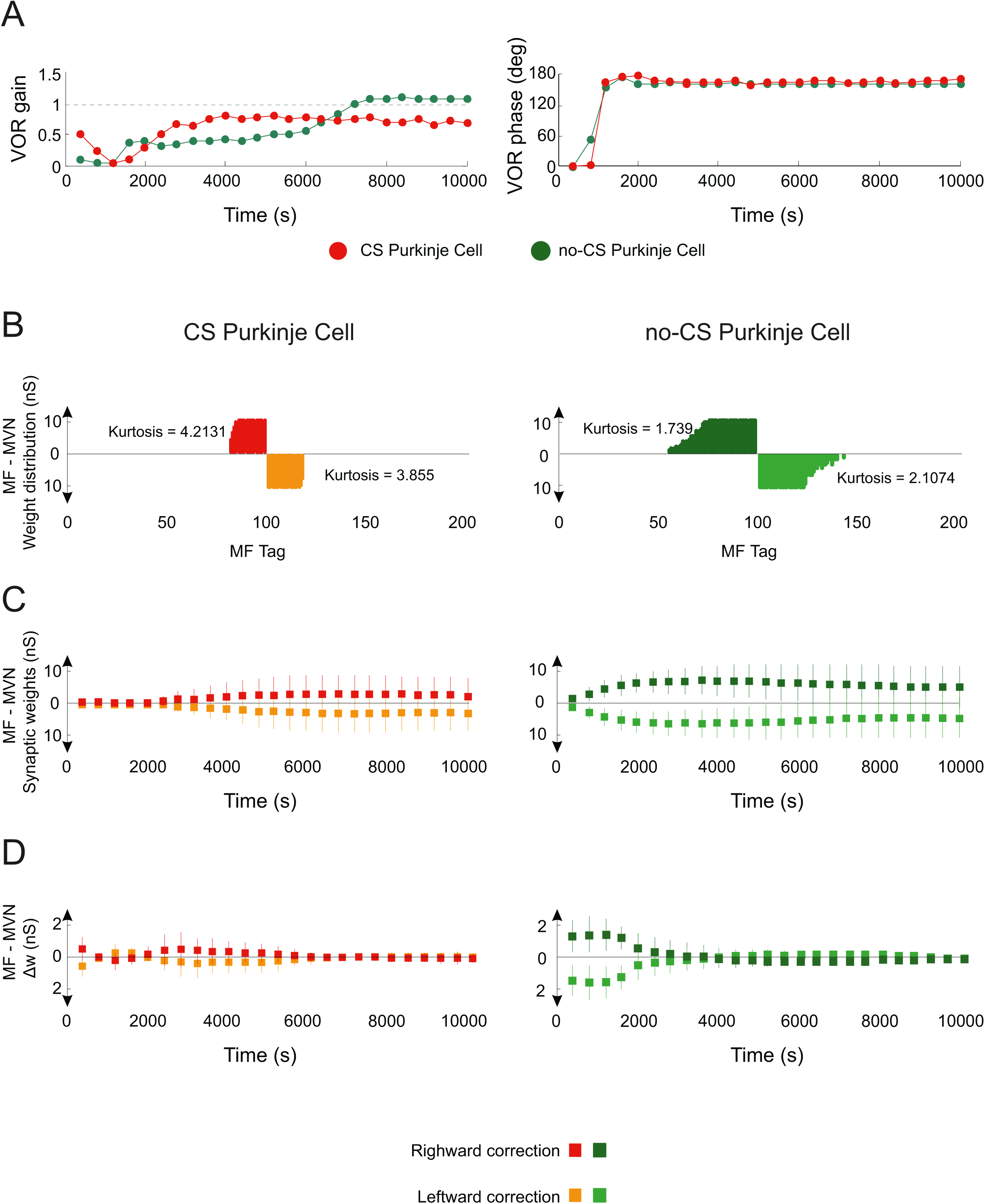
Role of Purkinje spike burst-pause dynamics in VOR cerebellar adaptation. **(A)** VOR gain and phase adaptation with (purple curve) and without (green curve) CF-evoked Purkinje spike burst-pause dynamics. VOR cerebellar adaptation starts with zero gain owing to the initial synaptic weights at PF and MVN afferents (Table 5). Purkinje spike burst-pause dynamics provides better VOR gain adaptation (in terms of both rate and precision) converging to values within [0.8-0.9], which is consistent with experimental data [36, 41, 42]. **(B)** Purkinje complex spiking allows a sparser weight distribution (with higher Kurtosis) to be learnt at MF-MVN synapses, with significantly lesser MF afferents needed for learning consolidation. **(C)** The model endowed with Purkinje complex spiking updates less MF afferents during learning consolidation but their synaptic range is fully exploited. **(D)** The averaged synaptic weight variations are more selective during the adaptive process in the presence of Purkinje spike burst-pause dynamics, yet the standard deviation remains equal.

A more accurate VOR gain adaptation in the presence of Purkinje complex spiking reflected a more selective synaptic modulation across learning (Figs 4B-D). In particular, Purkinje spike burst-pause dynamics facilitated a sparser weight distribution at MF-MVN synapses (Fig 4B), which ultimately shaped VOR adaptation [18]. Indeed, Purkinje burst sizes reflected the sensed errors [14], thus regulating the inhibitory action of Purkinje cells on MVN, and inducing error-dependent LTD at MF-MVN synapses (see Materials & Methods). On the other hand, post-complex spike pauses (disinhibiting MVN) induced error-dependent LTP at MF-MVN synapses (the larger the error, the larger the burst size, and the wider the post-complex spike pause, Fig 2B). At the beginning of VOR adaptation, the error was larger, and so were the burst and pause durations. Because the durations of pauses remained always larger than bursts (Fig 2B), LTP dominated over LTD at MF-MVN synapses, increasing the learning rate. Therefore, the spike burst-pause dynamics enhanced the precision of cerebellar adaptation at MVN cells, by *(i)* recruiting the strictly necessary MF-MVN projections (i.e., higher kurtosis value of the synaptic weight distribution, Fig 4B), *(ii)* making a better use of the synaptic range of selected projections (larger standard deviations with lower overall gains; Fig 4C), and the rate by *(iii)* varying synaptic weights selectively (lower averaged synaptic weight variations; Fig 4D).

### Purkinje spike burst-pause dynamics facilitates VOR phase-reversal learning

Phase-reversal VOR is induced when a visual stimulus is given simultaneously in phase to the vestibular stimulation but at greater amplitude (10% more) [25]. This creates a mismatch between visual and vestibular stimulation making retinal slips to reverse direction[43]. Cerebellar learning is deeply affected by VOR phase reversal since the synaptic weight distribution at both PF-Purkinje cell and MF-MVN synapses must be reversed. Here, we first simulated an h-VOR adaptation protocol (1 Hz) during 10000 s (as before). Then, h-VOR phase reversal took place during the next 12000 s. Finally, the normal h-VOR had to be restored during the last 12000 s (Fig 5). Our results suggest that Purkinje spike burst-pause dynamics were instrumental to phase-reversal VOR gain adaptation (Fig 5A), allowing for fast VOR learning reversibility consistently with experimental recordings [3] (Fig 5B). Conversely, the absence of Purkinje complex spiking led to impaired VOR phase-reversal learning with significant interference (Figs 5A,B). The two models (i.e., with and without Purkinje complex spiking) behaved similarly in terms of VOR phase adaptation during the same reversal learning protocol (S5-1 Fig).

**Figure 5.**
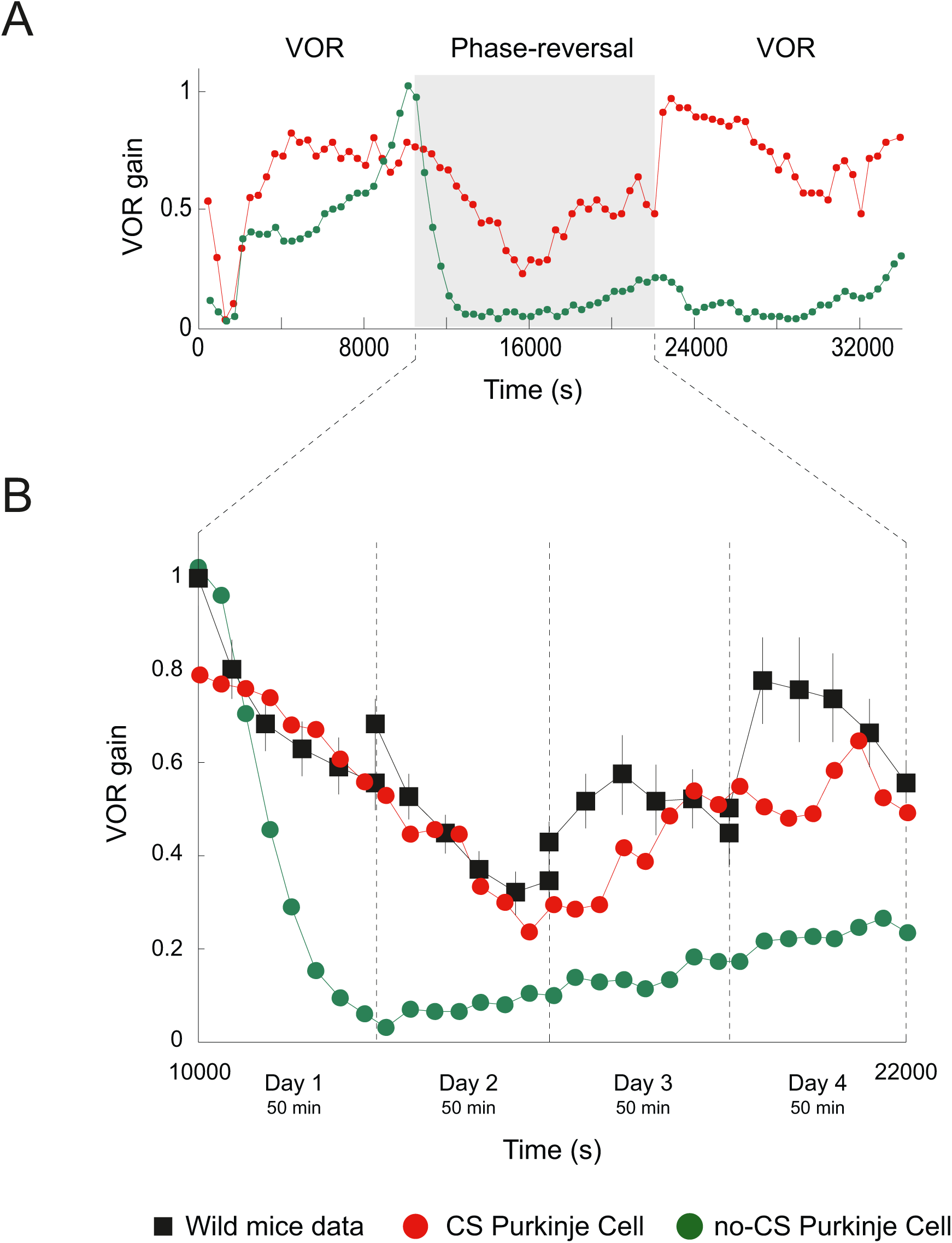
Purkinje spike burst-pause dynamics facilitates VOR phase-reversal learning. **(A)** VOR gain adaptation with (red curve) and without (green curve) Purkinje spike burst-pause dynamics during: VOR adaptation (first 10000 s), phase-reversal learning (subsequent 12000 s), and normal VOR restoration (remaining 12000 s). **(B)** Purkinje spike burst-pause dynamics provides fast learning reversibility, consistently with experimental recordings [3]. By contrast, phase-reversal VOR learning is impaired in the absence of Purkinje complex spiking. See S5-1 Fig for the time course of VOR phase-reversal learning.

VOR phase-reversal learning demanded first the reduction of the VOR gain, which can be regarded as a ‘forgetting phase’ (Fig 5B, days 1&2). Then, a ‘synchronisation phase’ took place with a reverse adaptive action that gradually increased the VOR gain (Fig 5B, days 3&4). During the forgetting phase, LTD dominated over LTP at MF-MVN synapses (Purkinje burst sizes were maximal), thus erasing the memorised weight patterns. During the synchronisation phase, Purkinje post-complex spike pauses led to a dominant LTP at MF-MVN synapses, reversing the learnt configuration. The interplay between bursts and post-complex spike pauses allowed synaptic adaptation at MF-MVN projections to be highly selective, which resulted in a sparser weight distribution as compared to the case without Purkinje complex spiking (Fig 6A). Therefore, VOR reverse learning required the adjustment of fewer MF-MVN synapses, thus facilitating the eye counteraction of the head velocity movement (S6-1 Fig), and the weight distribution was reshaped more efficiently with negligible interferences from the previously learnt patterns (Figs 6B, C).

**Figure 6.**
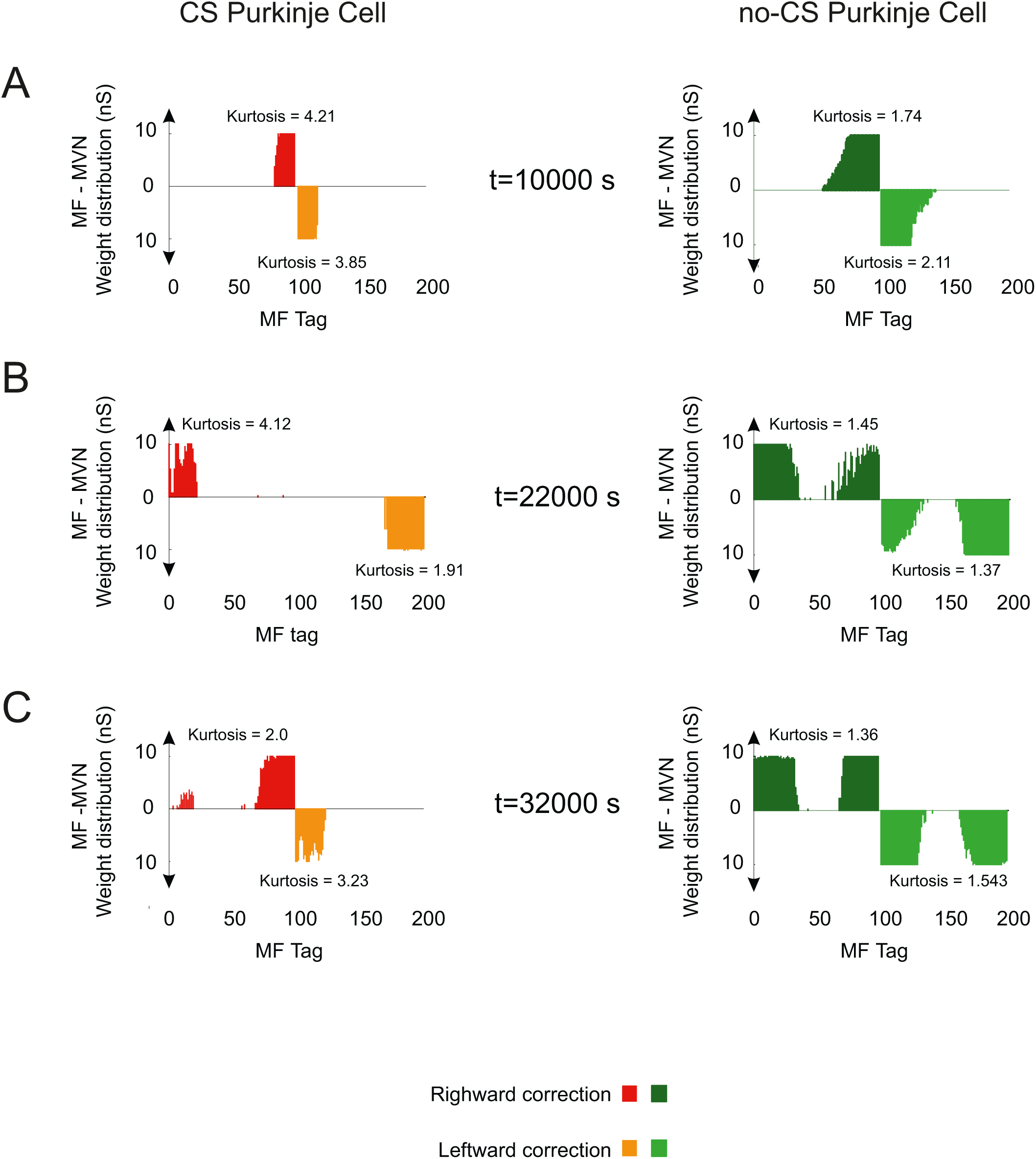
Evolution of synaptic weight distributions during VOR phase-reversal learning. (**A**) Only the sparser and more selective distribution of MF-MVN synaptic weights resulting from the interplay between bursts and post-complex spike pauses facilitates an efficient reshaping of the learnt patterns (**B**), allowing phase-reversal learning to be achieved (**C**).

### LTP blockades (by dominant LTD) during REMs explain reversal VOR gain discontinuities between training sessions

VOR phase-reversal learning can take place across several days [3] (Fig 5). Dark periods in-between training sessions cause reversal VOR gain discontinuities (Fig 7). This phenomenon has been assumed to result from the decaying of synaptic weights back to their initial values during sleep [3]. However, the mechanisms underlying this decaying process remain unknown. We explored possible cerebellar LTD/LTP balance modulation scenarios occurring during sleep as a consequence of changes in cerebellar activity. During rapid eye movement sleep (REMs), the mean firing activity of Purkinje cells shows increased tonic firing and decreased bursting in both frequency and size [44]. The CF average activity during REMs remains constant at a low frequency regime, showing a tendency in many IO neurons to diminish their overall frequency [45]. The activation of MFs varies during REMs, unrelatedly to any apparent behavioural changes, up to 60 MF/s on average [45].

**Figure 7.**
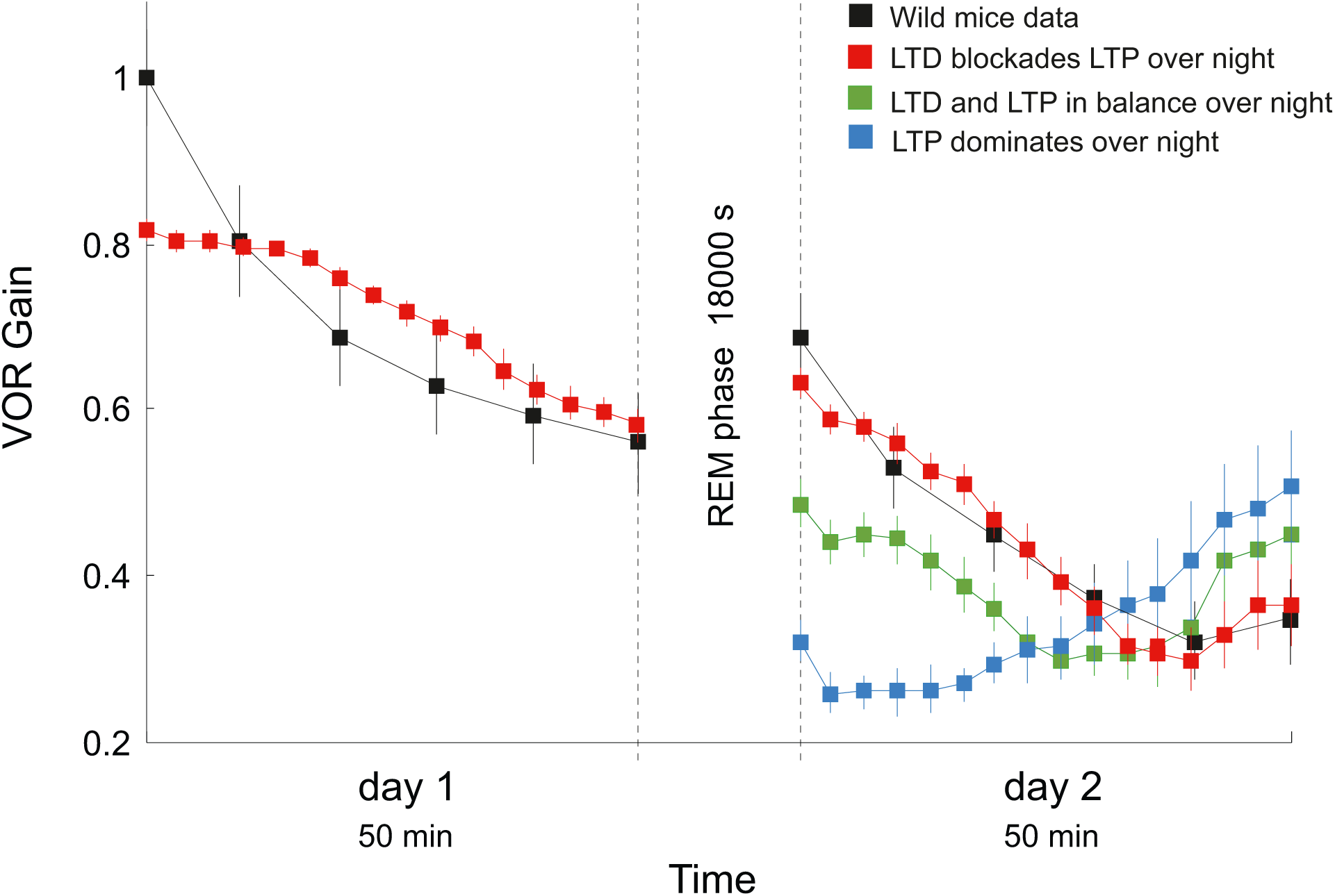
LTP blockades (due to dominant LTD) during REMs explain reversal VOR gain discontinuities between training sessions. We simulated 6 REMs stages (for a total of 18000 s of simulation) between day 1 and 2 of VOR phase-reversal learning. High levels of MF activity (10 Hz) leads to a dominance of LTP at both PF-Purkinje cell and MF-MVN synapses during REMs. Hence, during REMs the cerebellar model keeps ‘forgetting’ the memory traces as during day 1 (blue curve). A smaller MF activity (2.5 Hz) leads to a balance of LTP (driven by vestibular activity) and LTD (driven by the CFs). Thus, the model tends to maintain the synaptic weights learnt during day 1(orange curve). A very low MF activity (1 Hz) makes LTD to block LTP at PF-Purkinje and MF-MVN synapses. Under this third hypothesis, the synaptic weights tend to decay back towards their initial value (purple curve) in accordance with experimental data [3]. See S7-1 Fig for the modelled probabilistic Poisson process underpinning CF activation.

We modelled Purkinje cell, CF and MF activities during REMs. CFs were stochastically activated at 1 Hz [44, 45] following a Poisson distribution (S7-1 Fig). CF activations were also modulated to generate a large event in the Purkinje soma able to elicit bursts of 3 spikes on average [44]. MFs were stochastically activated by mimicking their activity during REMs (with an upper bound firing rate of 8-13 Hz). We tested three hypotheses, based on different levels of cerebellar activity during 6 REMs stages of 3000 s each (i.e., 18000 s of simulation) between days 1 and 2. In the first scenario, we considered high levels of MF activity (average firing rate 10 Hz), which led to a dominance of LTP at both PF-Purkinje cell and MF-MVN synapses during REMs. Consequently, the cerebellar model kept ‘forgetting’ the memory traces as during the reversal VOR learning of day 1 (Fig 7, blue curve). In the second scenario, we considered an average MF activity of 2.5 Hz, which made the LTP driven by vestibular activity to counterbalance the LTD driven by the CFs. Under this condition, the cerebellar model consolidated reversal VOR adaptation thus maintaining the synaptic weights at PF-Purkinje and MF-MVN synapses (Fig 7, green curve). Finally, we considered a low level of MF activity (average 1 Hz), which made LTD to block the LTP action driven by the vestibular (MF) activity. Under this third scenario, the cerebellar model showed a consistent tendency for weights at PF-Purkinje and MF-MVN synapses to decay back towards their initial value (Fig 7, red curve). Therefore, the model predicts that LTP blockade during REMs stages might underlie the reversal VOR gain discontinuities in-between training sessions, in agreement with experimental data [3] (Fig 7, black curve).

### Purkinje complex spike-pause dynamics under stationary VOR conditions

During transient VOR adaptation and phase reversal learning, retina slips were large causing vigorous CF discharges (up to 10 Hz) to encode the sensed errors. Consequently, Purkinje cell complex spike-pauses were elicited at high frequency during adaptation (Fig 8A). As the VOR error decreased, the frequency of CF-evoked Purkinje bursts decayed to ∼1 Hz upon completion of adaptation (Fig 8B). Therefore, during post (and pre) VOR adaptation, model Purkinje tonic Na^+^ spike output dominated and Purkinje cells tended to fire steadily (similar to spontaneous activity) with only rare complex spike-pause firing. Under stationary VOR conditions, (i.e., during pre/post VOR adaptation) model CFs were stochastically activated at ∼1 Hz (S7-1 Fig shows the Poisson-based generative model for the IO firing). Such a CF baseline discharge at ∼1 Hz allowed non-supervised LTP to be counterbalanced at PF-Purkinje cell synapses (see Materials & Methods), thus preserving pre/post cerebellar adaptation.

**Figure 8.**
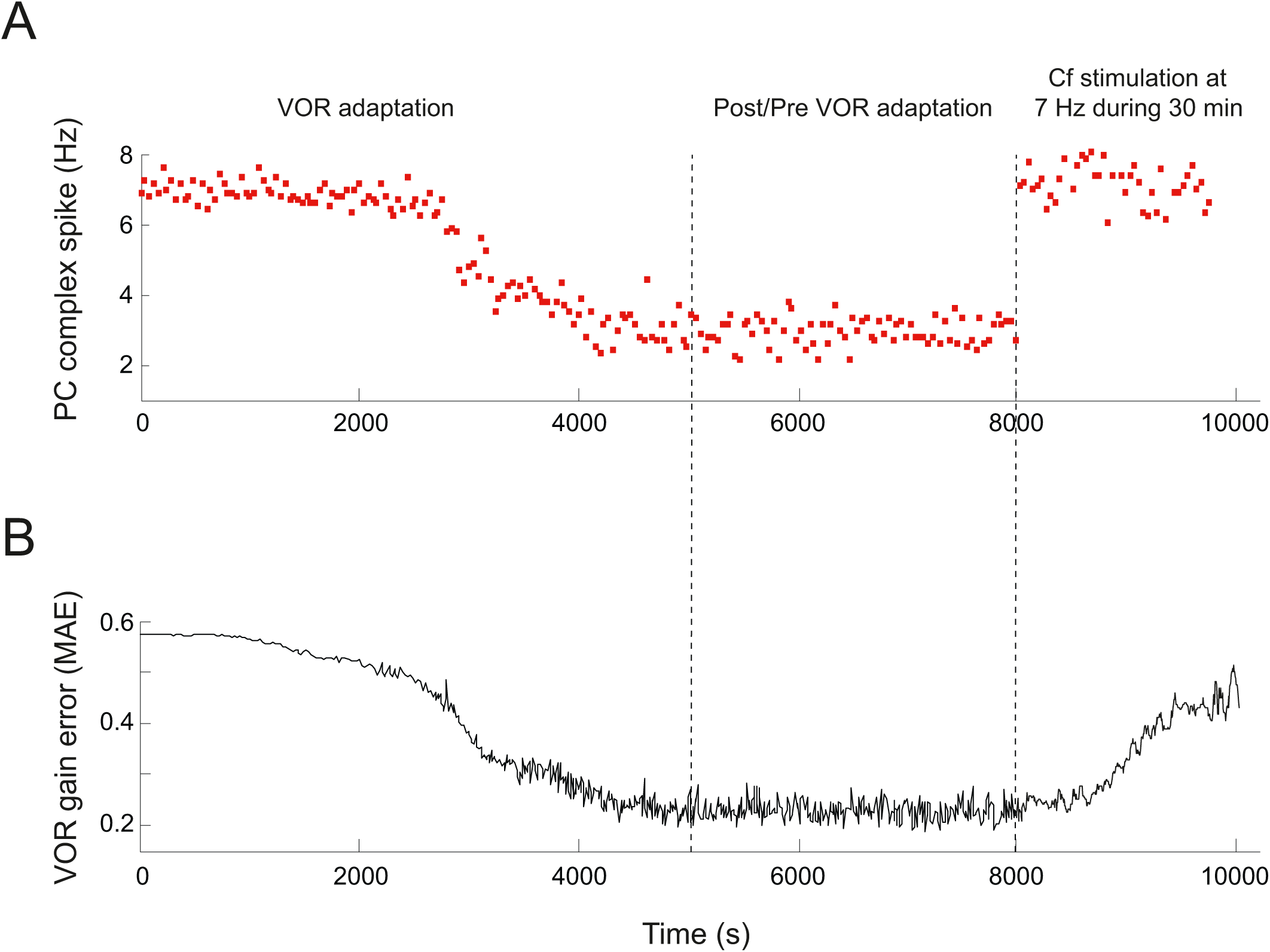
Purkinje complex spike-pause frequency and VOR gain error during adaptation and post/pre adaptation. **(A)** The frequency of Purkinje complex spike-pauses diminishes through VOR adaptation from 8-9 Hz to 1-2 Hz under a sinusoidal vestibular stimulus of ∼1 Hz. After VOR adaptation, a direct random stimulation of CFs at 7 Hz during 30 min as in [46] impairs the VOR reflex. **(B)** Evolution of the VOR gain error (Mean Absolute Error) during adaptation, post-adaptation, and artificial random stimulation of CFs.

Luebke and Robinson [46] found that directly stimulating CFs at 7 Hz during 30 min after 3 days of VOR adaptation would impair the reflex. Model CFs discharged at frequencies larger than 1 Hz only to signal retina slips (i.e., during VOR adaptation). However, a direct (and error independent) high-frequency stochastic stimulation of CFs would lead to VOR impairment. To illustrate this, we simulated a protocol similar to the one used by [46]. As expected, the number of CF-evoked Purkinje burst-pauses increased as the CF frequency was artificially incremented through a 7 Hz direct stimulation (Fig 8A).Therefore, the VOR gain error tended to increase indicating an impairment/blockade of the acquired reflex (Fig 8B) and a decrease in VOR gain even with similar CFs discharges observed during VOR adaptation.

## Discussion

Marr and Albus theory [47, 48] elicited a large body of research on the link between the cellular and network properties of the cerebellum and behavioural adaptation. This extensive effort crystallised into a broad range of cerebellar models based on divergent premises. On the one hand, detailed models were grounded on cellular and synaptic properties observed experimentally [49–54]. Most of these biophysical models did not aim at driving behavioural adaptation explicitly through network-level dynamics. On the other hand, numerous large-scale solutions were engineered to be computationally efficient for learning sensorimotor tasks, regardless of the anatomo-functional constraints governing cellular and network cerebellar processes [55–58]. The approach presented here conjugates these two vantage points and focuses on the role of the multiple spiking patterns of Purkinje cells in cerebellar adaptation. It is well known that Purkinje cells can express fast tonic firing as well as a characteristic burst-pause spiking pattern in response to excitatory parallel fibre (PF) and climbing fibre (CF) inputs [40]. Nevertheless, we address the still uncovered question of how these different spiking patterns regulate the inhibitory action of Purkinje cells onto targeted medial vestibular nuclei (MVN) and ultimately shape the adaptive behavioural control mediated by the cerebellum.

We model cerebellar-dependent adaptation of the rotational vestibulo-ocular reflex (VOR) (Fig 1A). For natural head rotation frequencies (0.5–5.0 Hz), the VOR gain (i.e., eye velocity divided by head velocity) and the VOR phase shift (i.e., the time lag between eye and velocity profiles) are close to 1 and 180°, respectively [7]. Thus, synchronised counter-phased eye and head movements stabilise visual targets on the fovea, minimising retina slips and improving visual acuity [59]. Cerebellar learning, and particularly Purkinje cell response adaptation, is necessary to mediate online changes in VOR gain control [60, 61]. The cerebellar model presented here mimics the main properties of the cerebellar microcircuit, and it embodies spike-based LTP/LTD plasticity mechanisms at multiple synaptic sites (Fig 1C). At the core of the spiking cerebellar network, a detailed single-compartment model of Purkinje cell reproduces the characteristic tonic, complex spike, and post-complex spike pause patterns [62, 63]. In order to focus on how CF-evoked spike burst-pause dynamics of Purkinje cell responses can regulate the adaptive output of the cerebellum, we also use a Purkinje neuron model that cannot express complex spike firing (i.e., it can only operate in tonic mode). The main finding of this study is that the CF-evoked spike burst-pause dynamics of the Purkinje cell is a key feature for supporting both early and consolidated VOR learning. The model predicts that properly timed and sized Purkinje spike burst-pause sequences are critical to: (1) gating the contingent association between vestibular inputs (about head rotational velocity) and MVN motor outputs (to determine counter-rotational eye movements), mediating an otherwise impaired VOR coarse acquisition; (2) allowing the LTD/LTP balance at MF-MVN synapses to be accurately shaped for optimal VOR consolidation; (3) reshaping previously learnt synaptic efficacy distributions for VOR phase-reversal adaptation. Finally, the model predicts that the reversal VOR gain discontinuities observed after sleeping periods in-between training sessions [3] are due to LTD/LTP balance modulations (and in particular LTP blockades) occurring during REM sleep as a consequence of changes in cerebellar activity.

This work assumes a gradually modulated CF activity capable of instructing a ‘teaching’ signal to Purkinje cells [64]. The type of information conveyed by CFs onto Purkinje cells (and its potential role in sensorimotor adaptation) is under debate. On the one hand, CFs have been hypothesised to carry a binary feedback-error signal computed by IO [65]. On the other hand, recent studies have questioned the hypothesis of a binary CF signal by demonstrating that the duration of Purkinje cell complex spikes (evoked by CF afferents) can be accurately adjusted based on information that a binary teaching signal could not support [14, 15, 66–68]. Our model embraces this second hypothesis. It must also be noted that the overall assumption about IO-mediated feedback-error learning has been contrasted by a body of research that focused on the periodic nature of CF activity. These works put the CF signalling in relation to the timing aspects of motion [69, 70] and, in particular, to the onset of motion [71]. The controversy about the nature of CF activity has been further roused by the fact that IO functional properties have so far not been univocally identified [60, 72–74].

The model presented here captures the fact that similar CF discharges occur during both VOR gain increase and decrease adaptation [75, 76]. CFs encode the retinal slips that drive VOR adaptation [77]. The direction of retinal slips relative to the vestibular stimulus induces either an increase or a decrease in VOR gain [78]. Interestingly, the relation between CF activity and the induction of plasticity at Purkinje cell synapses is described as a gating mechanism that varies under these two VOR adaptation paradigms [76]. Furthermore, optogenetic CF stimulation in VOR gain-decrease paradigms suggest that changes in Purkinje cell complex spike responses do not only depend upon CF activation [76]. Our cerebellar model accounts for these observations by means of the mechanism that balances LTD/LTP plasticity at PF-Purkinje cell synapses. During VOR *gain–increase* adaptation, LTD predominantly blocks LTP at modelled PF-Purkinje cell synapses. This results in a synaptic efficacy decrease as a CF spike reaches the target Purkinje cell (error-related signal). In particular, a CF spike is more likely to depress a PF-Purkinje cell synapse if the PF has been active within 50-150 ms of the CF spike arrival [79–81]. Increasing LTD at PF-Purkinje cell synapses reduces the inhibitory action of Purkinje cells on MVN activity, which in turn increases the VOR gain. During VOR *gain–decrease* adaptation [25, 75], LTP dominates at PF–Purkinje cell synapses, despite the fact that CF inputs are similar to those occurring during gain-increase phases. A raise in synaptic efficacy at PF-Purkinje cell synapses increases the inhibition of MVN neurons, which in turn reduces the VOR gain. LTP at modelled PF-Purkinje cell synapses is non-supervised and it strengthens a connection upon each PF spike arrival at the target Purkinje cell. This plasticity mechanism does not need to modulate the input provided by CFs (and then the CF-evoked spike burst-pause dynamics of Purkinje cells) to counter LTD and decrease the VOR gain, in accordance to in-vitro experiments [82- 84].

The model suggests that CF-evoked Purkinje cell spike burst-pause dynamics are critical to shape MF-MVN synapses, as to optimise the accuracy and consolidation rate of VOR adaptation. We show that burst and spike pause sequences facilitate sparser MF-MVN connections, which increases coding specificity during the adaptation process. The results predict that the spike burst-pause dynamics should be central to retune MF-MVN synapses during VOR phase-reversal adaptation. First, it is shown that blocking complex spike responses (and post-complex spike pauses) in Purkinje cells impairs reverse VOR adaptation. More strikingly, the results indicate that Purkinje cell bursting and spike pauses ensure the reversibility of the adaptation process at MF-MVN synapses. Bursts selectively facilitate LTD at MF-MVN connections, which rapidly erases previously learnt memory traces at these synapses. Subsequently, post-complex spike pauses induce strong LTP at MF-MVN synapses, which allows the cerebellar output to become rapidly reverse-correlated to the sensed error. In addition, the memory consolidation of VOR adaptation during sleeping [3, 85, 86] is also supported by the CF-evoked Purkinje cell spike burst-pause dynamics. CF stochastically activations at a low frequency (0.9 Hz) during REMs stages maintain abase Purkinje bursting that ultimately facilitates LTP blockades at PF-Purkinje cell and MF-MVN synapses, and it preserves the on-going learning process.

The cerebellar model endowed with CF-evoked Purkinje cell spike burst-pause dynamics performs better, in terms of adaptation accuracy and consolidation rate, than a model with Purkinje cells expressing tonic firing only. CF-evoked spike burst-pause patterns appear particularly useful in a disruptive task such as VOR phase-reversal adaptation. Nevertheless, our results indicate that complex spikes, post-complex spike pauses, and their relative modulation, are not essential for VOR control learning and adaptation. This is in agreement with recent experimental findings challenging the hypothesis that Purkinje cell complex spikes are necessarily required in cerebellar adaptation, and suggesting that their role in motor learning is paradigm dependent [74, 87]. Overall, this work provides insights on how the signals provided by the CFs may instruct, either directly or indirectly, plasticity at different cerebellar synaptic sites [64]. The results point towards a key role of CF-evoked Purkinje cell spike burst-pause dynamics in driving adaptation at downstream neural stages. This testable prediction may help to better understanding the cellular-to-network principles underlying cerebellar-dependent sensorimotor adaptation.

## Materials & Methods

### VOR Analysis and Assessment

We simulated horizontal VOR (h-VOR) experiments with mice undertaking sinusoidal (∼1 Hz) whole body rotations in the dark [36]. The periodic functions representing eye and head velocities (Fig 1A) were analysed through a discrete time Fourier transform. The **VOR gain** was calculated as the ratio between the first harmonic amplitudes of the forward Fourier eye– and head–velocity transforms:

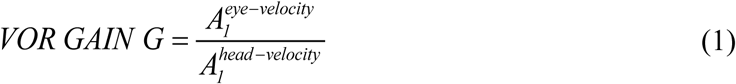

In order to assess the **VOR shift phase**, the cross-correlation of the eye and head velocity time series was computed:

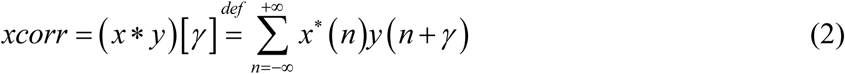

where *x^*^* is the complex conjugate of *x,* and *γ* the lag (i.e. shift phase). The ideal eye and head velocity lag is ±0.5 after normalisation, with cross-correlation values ranged within [−1, 1], which is equivalent to a phase shift interval of [−360° 360°].

### Cerebellar Spiking Neural Network Model

The cerebellar circuit was modelled as a feed–forward loop capable of compensating head movements by producing contralateral eye movements (Fig 1B). The connectivity and the topology of the simulated cerebellar network involved five neural populations: mossy fibres (MFs), granule cells (GCs), medial vestibular nuclei (MVN), Purkinje cells, and inferior olive (IO) cells [29, 88–91]. During simulated 1 Hz head rotations, sensorimotor activity was translated into MF activity patterns that encoded head velocity. MFs transmitted this information to both MVN and GCs. The latter generated a sparse representation of head velocity signals, which was sent to Purkinje cells through the PFs. Purkinje cells were also driven by the CFs, which conveyed the teaching signal encoding sensory error information (i.e., retina slips due to the difference between actual and target eye movements, [77]). Finally, Purkinje cells’ output inhibited MVN neurons, which closed the loop by shaping cerebellar-dependent VOR control. The CF-Purkinje cell-MVN subcircuit was divided in two symmetric micro-complexes for left and right h-VOR, respectively. The input-output function of the cerebellar network model was made adaptive through spike-timing dependent plasticity (STDP) at stake at multiple sites (Fig 1C). These STDP mechanisms led to both long-term potentiation (LTP) and long-term depression (LTD) of the ∼50000 synapses of the cerebellar model see [92]. This spiking neural network model was implemented in EDLUT [81, 93, 94]an efficient open source simulator mainly oriented to real time simulations.

#### Purkinje cell model

We considered a detailed Purkinje cell model [62, 63] consisting of a single compartment with five ionic currents:

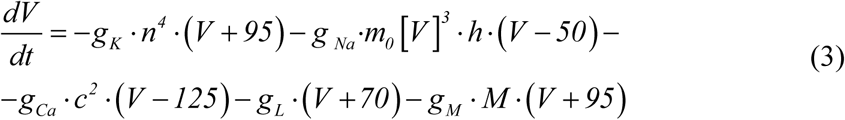

with *g_K_* denoting a delayed rectifier potassium current, *g _Na_* a transient inactivating sodium current, *g_Ca_* a high-threshold non-inactivating calcium current, *g_L_* a leak current, and *g_M_* a muscarinic receptor suppressed potassium current (see Table 1).

**Table 1.** Ionic conductance densities

| *Conductance type* | *Soma (mho/cm2)* |
| --- | --- |
| $g_K$—delayed rectifier potassium current | 0.01 |
| $g_{Na}$—transient inactivating sodium current | 0.125 |
| $g_{Ca}$—high threshold | 0.001 |
| $g_M$—muscarinic receptor | 0.75 |
| $g_L$—leak current (anomalous rectifier) | 0.02 |

The dynamics of each gating variable evolved as follows:

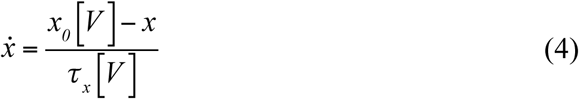

where x indicates the variables n, h, c, and M. The implemented equilibrium function is determined by the term *x_0_* [*V*] and time constant *τ_x_* [*V*] (Table 2).

**Table 2.** Ionic conductance kinetic parameters

| *Conductance type* | *Steady-state*<br>*Activation/Inactivation* | *Time constant (ms)* |
| --- | --- | --- |
| $g_K$—delayed rectifier<br>potassium current | $x_0[V] = \frac{1}{1 + e^{\frac{-V-29.5}{10}}}$ | $\tau_x[V] = \begin{cases} 0.25 + 4.35 \cdot e^{\frac{V+10}{10}} & \text{if } V \leq 10 \\ 0.25 + 4.35 \cdot e^{\frac{-V-10}{10}} & \text{if } V > 10 \end{cases}$ |
| $g_{Na}$—transient<br>inactivating sodium<br>current | $x_0[V] = \frac{1}{1 + e^{\frac{V-59.4}{10.7}}}$ | $\tau_x[V] = 0.15 + \frac{1.15}{1 + e^{\frac{V+33.5}{15}}}$ |
| $m_0[V]$ | $m_0[V] = \frac{1}{1 + e^{\frac{-V-48}{10}}} \cdot m$ | |

| | *Forward Rate Function* ($\alpha$) | *Backward Rate Function* ($\beta$) |
| --- | --- | --- |
| $g_{Ca}$-high threshold | $\alpha = \frac{1.6}{1 + e^{-0.0072 \cdot (V-5)}}$ | $\beta = \frac{0.02 \cdot (V + 8.9)}{e^{\frac{V+8.9}{5}}}$ |
| $g_M$-muscarinic<br>receptor suppressed<br>potassium current | $\alpha = \frac{0.3}{1 + e^{\frac{-V-2}{5}}}$ | $\beta = 0.001 \cdot e^{\frac{-V-70}{18}}$ |
|  | *Steady-state*<br>*Activation/Inactivation* | *Time constant(ms)* |
| | $x_0[V] = \frac{\alpha}{\alpha + \beta}$ | $\tau_x[V] = \frac{1}{\alpha + \beta}$ |

The sodium activation variable was replaced and approximated by its equilibrium function *m_0_* [*V*]. M-current presents a temporal evolution significantly slower than the rest of the five variables thus provoking a slow-fast system able to reproduce the characteristic Purkinje cell spiking modes (Fig 2).

The final voltage dynamics for the Purkinje [62, 63]cell model was given by:

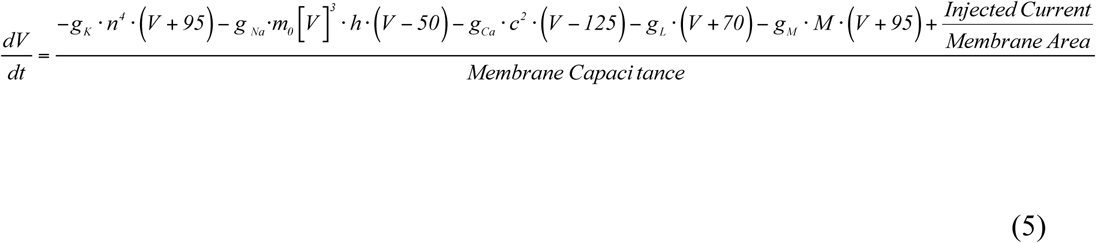

where the parameters *Membrane Area* and *Membrane Capacitance* are provided in Table 3, and *Injected Current* is the sum of all contributions received through individual synapses (see Eqs. 6–8 below).

**Table 3.** Geometrical parameters:

| *Geometrical parameters* |  |
| --- | --- |
| Cylinder length of the soma | 15 $\mu m$ |
| Radius of the soma | 8 $\mu m$ |
| Membrane Capacitance | 1 $\mu F/cm^2$ |
| Axial resistivity | 100 $\Omega-cm$ (axom) 250 $\Omega-cm$ (dendrites) |
| Number of segments | 1 |

First, we validated the detailed Purkinje cell model (Eqs. 3–5) in the Neuron simulator. Subsequently, we reduced the Purkinje cell model to make it compatible with an event-driven lookup table (EDLUT simulator https://github.com/EduardoRosLab/edlut) for fast spiking neural network simulation [81, 93]. In the reduced Purkinje cell model, *I_K_* and *I_Na_* currents were implemented through a simple threshold process that triggers the generation of a triangular voltage function each time the neuron fires [95]. This triangular voltage depolarisation drives the state of ion channels similarly to the original voltage depolarisation during the spike generation.

#### Other cerebellar neuron models

The other cerebellar neurons (granule cells, MVN cells,…) were simulated as leaky integrate–and–fire (LIF) neurons, with excitatory (AMPA) and inhibitory (GABA) chemical synapses:

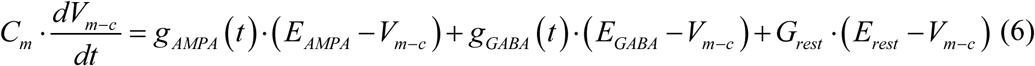

where *C_m_* denotes the membrane capacitance, *E_AMPA_* and *E_GABA_* are the reversal potential of each synaptic conductance, *E_rest_* is the resting potential, and *G_rest_* indicates the conductance responsible for the passive decay term towards the resting potential. Conductances *g_AMPA_* and *g_GABA_* integrate all the contributions received by each receptor type (AMPA and GABA) through individual synapses and they are defined as decaying exponential functions [81, 96]:

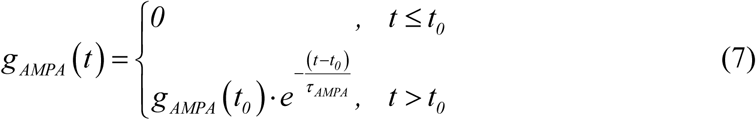

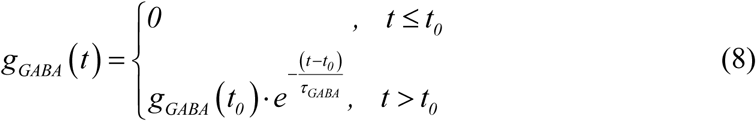

with *t* representing the simulation time, *t_0_* being the time arrival of an input spike, and *τ_AMPA_* and *τ_GABA_* denoting the decaying time constant for AMPA and GABA receptors, respectively.

Note that we also used the LIF neuronal model (Eqs. 6–8) to simulate Purkinje cells that could express tonic spike firing only (Fig 3B). These Purkinje cells without CF-evoked spike burst-pause dynamics provided a coarse phenomenological model reminiscent of Kv3.3-deficient Purkinje neurons (as in Kcnc3 mutants, in which the absence of voltage-gated potassium channel Kv3.3 compromises spikelet generation within complex spikes of cerebellar Purkinje cells) [97]. Table 4 summarises the parameters used for each cell and synaptic receptor type.

**Table 4.**
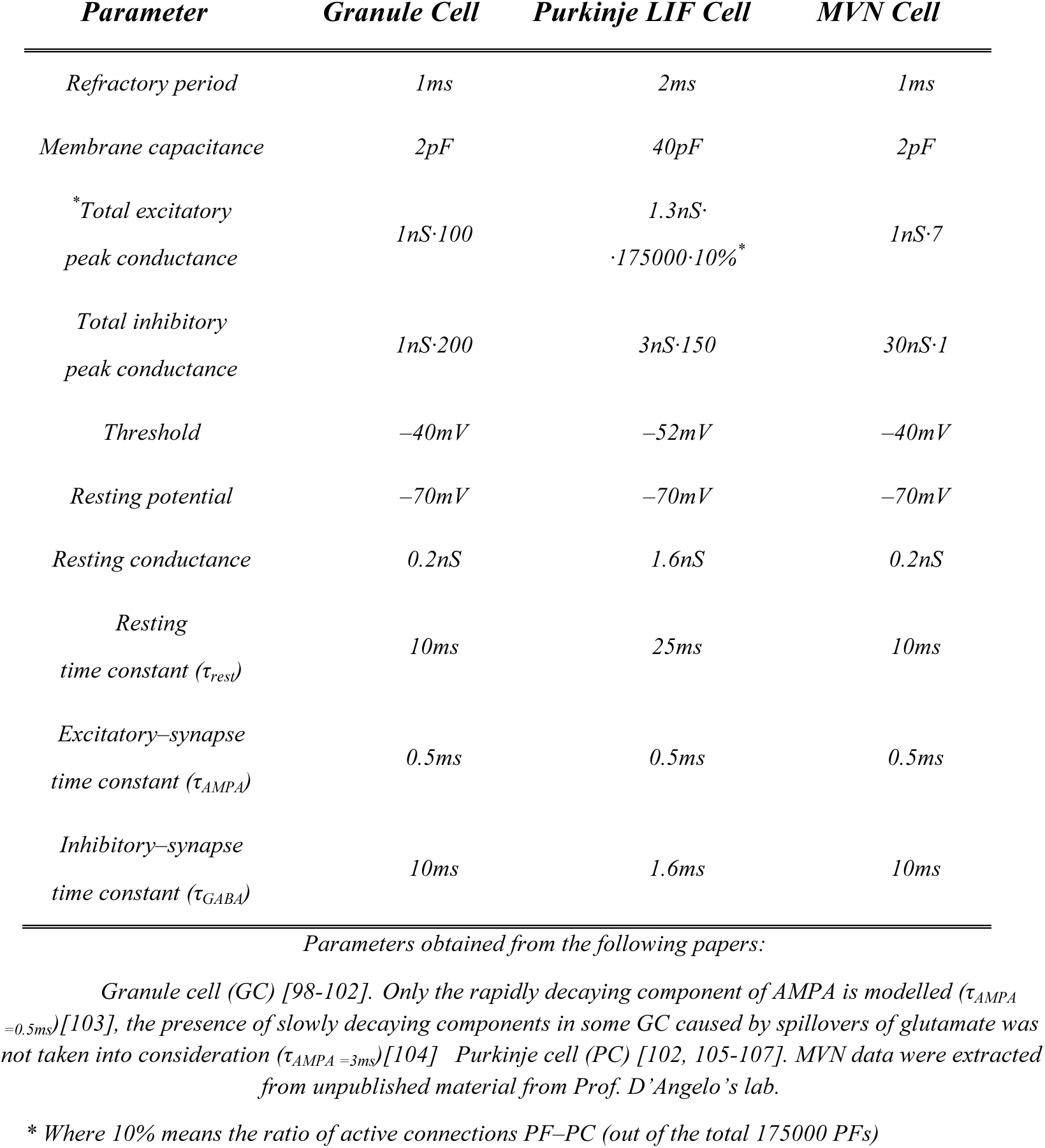
Parameters of the LIF cell types

#### Cerebellar neural population models

##### Mossy fibres (MFs)

N=100 MFs were modelled as LIF neurons (Eqs. 6–8). Consistently with the functional principles of VOR models of cerebellar control [3], the ensemble MF activity was generated following a sinusoidal shape (1 Hz with a step size of 0.002 ms) to encode head movements [3, 108, 109]. The overall MF activity was based on non-overlapping and equally sized neural subpopulations that allowed a constant firing rate of the ensemble MFs to be maintained over time. Importantly, two different times always corresponded to two different subgroups of active MFs ensuring to the overall constant activity. (Network connectivity parameters summarised in Table 5).

**Table 5.**
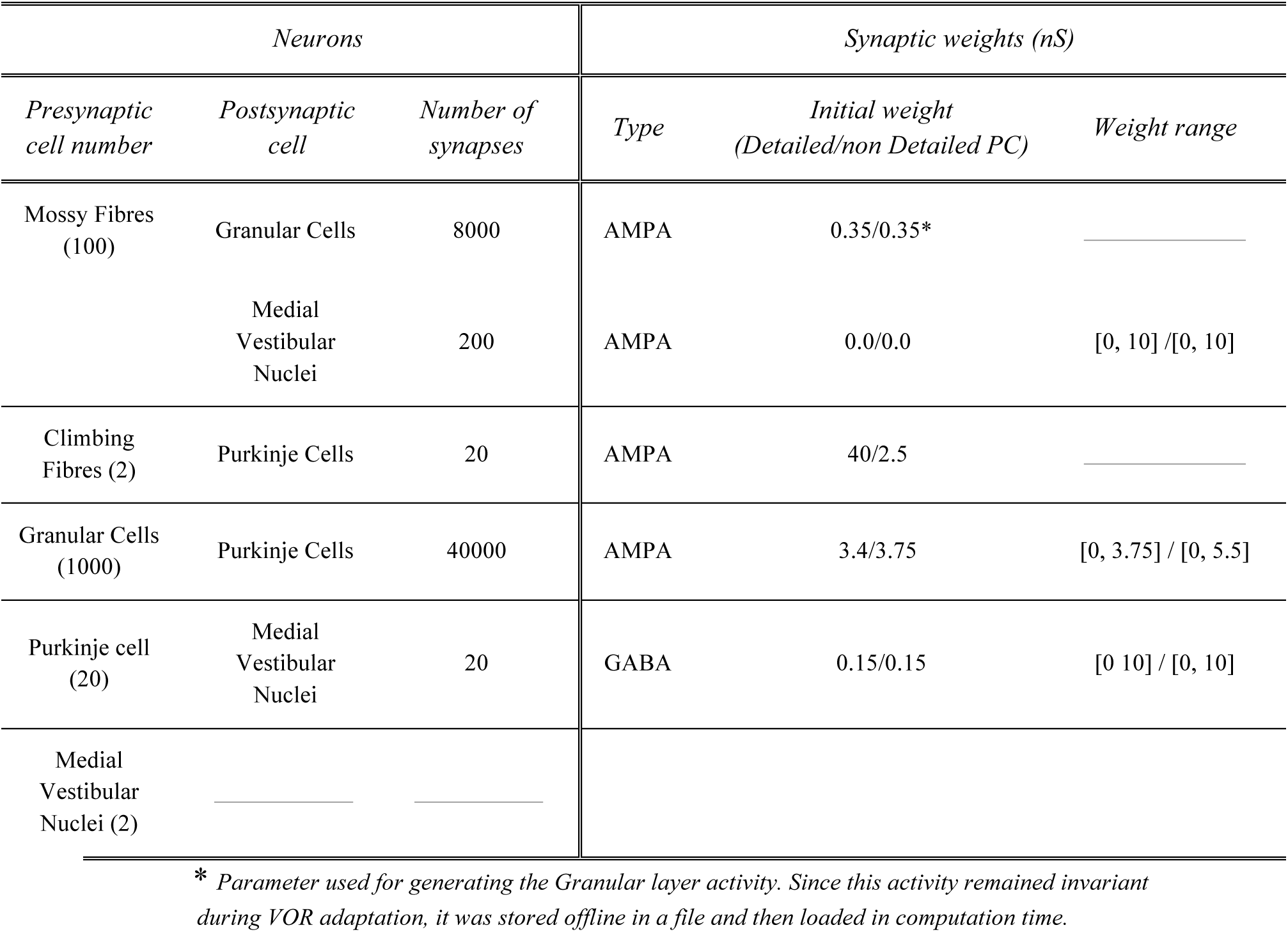
Summary of neurons and synapses.

##### Granular cells (GCs)

The granular layer included N=2000 GCs and it was implemented as a state generator [110–113], i.e. its inner dynamics produced time–evolving states even in the presence of a constant MF input [56]. The granular layer generated non-overlapped spatiotemporal patterns that were repeatedly activated in the same sequence during each learning trial (1 Hz rotation for 1 s)). 500 different states encoded each second of the 1 Hz learning trial, each state consisting of four non-recursively activated GCs.

##### Climbing fibres (CFs)

N=2 CFs carried the teaching signal (from the IO) to the population of Purkinje cells. The two CFs handled clockwise and counter–clockwise sensed errors. CF responses followed a probabilistic Poisson process. Given the normalised error signal *ε*(*t*) and a random number *η*(*t*) between 0 and 1, a CF fired a spike if *ε*(*t*) > *η*(*t*), otherwise it remained silent [79, 114, 115]. Thus, a single CF spike encoded well – timed information regarding the instantaneous error. Furthermore, the probabilistic spike sampling of the error ensured a proper representation of the whole error region over trials, while maintaining the CF activity below 10 Hz per fibre (similar to electrophysiological data; [116]. The evolution of the error could be sampled accurately even at such a low frequency [115, 117]. For the sake of computational efficiency, there are only 2 CFs (instead of 20 CFs). In the cerebellum, each PC is innervated by a single CF [118] coming from the associated IO at the olivary system. However, no olivary system is here considered and, consequently, CFs sensing clockwise and counter–clockwise errors are equally activated. It would suffice 1 CF sensing clockwise and 1 CF sensing anti-clockwise errors.

##### Purkinje cells

N=20 Purkinje cells were divided in two subpopulations of 10 neurons each. Each subpopulation received the inputs from one CF encoding the difference between (either rightward or leftward) eye and head movements. Each Purkinje cell also received 2000 PF inputs. Since real Purkinje cells are innervated by about 150000 PFs [119], the weights of the PF–Purkinje cells synapses of the model were scaled so as to obtain a biologically plausible amount of excitatory drive. Each of the two subgroups of 10 Purkinje cells targeted (through inhibitory projections) one MVN cell, responsible for either clockwise or counter-clockwise compensatory motor actions (ultimately driving the activity of agonist/antagonist ocular muscles).

##### Medial Vestibular Nuclei (MVN)

The activity of N=2 MVN cells produced the output of the cerebellar model. The two MVN neurons handled clockwise and counter– clockwise motor correction, respectively. Each MVN neuron received excitatory projections from all MFs (which determined the baseline MVN activity), and inhibitory afferents from the corresponding group of 10 Purkinje cells (i.e., the subcircuit IO– Purkinje cell–MVN was organised in a single microcomplex). MVN spike trains were translated into analogue output signals through a Finite Impulse Response filter (FIR) [120]. Let 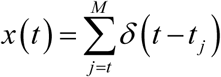 denote a MVN spike train, with *t_j_* being the firing times of the corresponding neuron. If *h(t)* indicates the FIR kernel, then the translated MVN output is:

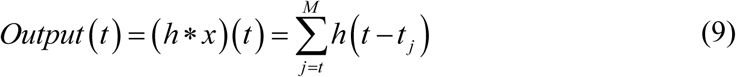

Note that a delay is introduced in the generated analogue signal. This delay is related to the number of filter coefficients and to the shape of the filter kernel *h(t)*. In order to mitigate this effect, we used an exponentially decaying kernel:

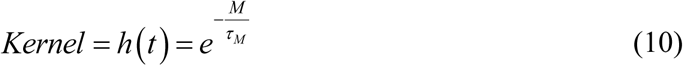

where *M* is the number of filter taps (one per integration step) and *τ_M_* is a decaying factor.

At each time step, the output signal value only depends on its previous value and on the input spikes in the same time step. Therefore, this filter is implemented by recursively updating the last value of the output signal. Importantly, this kernel is similar to postsynaptic current functions [121, 122], thus facilitating a biological interpretation. Furthermore, this FIR filter is equivalent to an integrative neuron [123].

#### Synaptic plasticity rules

##### PF–Purkinje cell synaptic plasticity

The LTD/LTP balance at PF–Purkinje cell synapses was based on the following rule (S3-1 Fig shows sensitivity analysis accounting for LTD/LTP balance):

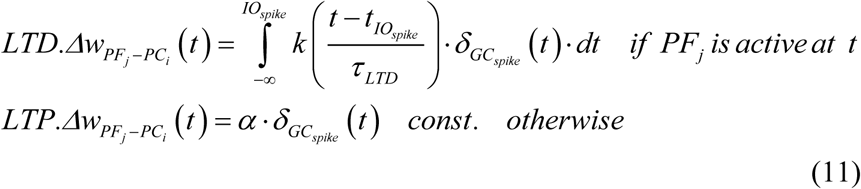

where Δ*W_PFj–PCi_(t)* denotes the weight change between the *j^th^* PF and the target *i^th^* Purkinje cell; *τ_LTD_* is the time constant that compensates for the sensorimotor delay (100ms); *δ_GR_* is the Dirac delta function corresponding to an afferent spike from a PF (i.e., emitted by a GC); and the kernel function *k(x)* is defined as [92]:

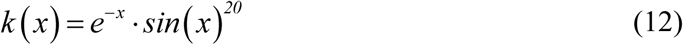

The convolution in Eq. 11 was computed on presynaptic PF spikes arriving 100 ms before a CF spike arrival, accounting for the sensorimotor pathway delay [65, 114, 115, 124]. Note that the kernel *k(x)* allows the computation to be run on an event–driven simulation scheme as EDLUT [81, 114, 115, 124], which avoids integrating the whole kernel upon each new spike arrival. Finally, as shown in Eq. 11, the amount of LTP at PF–Purkinje cell synapses was fixed, with an increase in synaptic efficacy equal to α each time a spike arrived through a PF to the targeted Purkinje cell.

##### MF–MVNsynaptic plasticity

The LTD/LTP dynamics at MF-MVN synapses was taken as (Fig. 3-1 shows sensitivity analysis accounting for LTD/LTP balance):

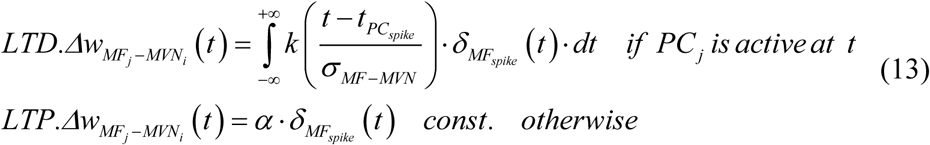

with Δ*W_MFj–MVNi(t)_* denoting the weight change between the *j^th^* MF and the target *i^th^* MVN. *σ_MF_*_-*DCN*_ standing for the temporal width of the kernel; *δ_MF_* representing the Dirac delta function that defines a MF spike; and the integrative kernel function *k(x)* defined as [92]:

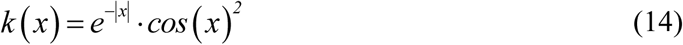

Note that there is no need to compensate the sensorimotor pathway delay at this site because it is already done at PF-Purkinje cell synapses (*τ_LTD_* in Eq. 11).

The STDP rule defined by Eq. 13 produces a synaptic efficacy decrease (LTD) when a spike from the Purkinje cell reaches the targeted MVN neuron. The amount of synaptic decrement (LTD) depends on the activity arrived through the MFs. This activity is convolved with the integrative kernel defined in Eq. (14). This LTD mechanism considers those MF spikes that arrive after/before the Purkinje cell spike arrival within the time window defined by the kernel. The amount of LTP at MF-MVN synapses is fixed (Ito, 1982;[92, 125], with an increase in synaptic efficacy each time a spike arrives through a MF to the targeted MVN.

##### Purkinje cell–MVNsynaptic plasticity

The STDP mechanism implemented at Purkinje cell-MVN synapses [92] consists of a traditional asymmetric Hebbian kernel

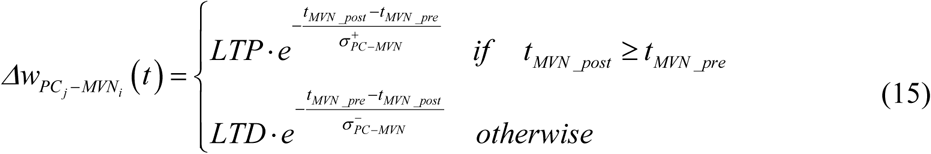

where Δ*W_Pqj–MVNi(t)_* is the weight change between the *j^th^* PC and the target *i^th^* MVN, 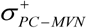 and 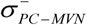 are the time constants of the depression potentiation and components set to 5ms and 15ms respectively; and LTD_max/_LTP_max_ (0.005/0.005) are the maximum weight depression/potentiation change per simulation step. The t_mvn_post_ and t_mvn_pre_ indicate the postsynaptic and presynaptic MVN spike time. This STDP rule is consistent with the fact that plasticity at Purkinje cell-MVN synapses depends on the intensity of MVN and Purkinje cell activities [20–23] and it provides a homeostatic mechanism in balancing the excitatory and inhibitory cell inputs to MVN [90, 126]. The source code is available at URL: http://www.ugr.es/∼nluque/restringido/CODE.rar (user: REVIEWER, password: REVIEWER).

## Acknowledgements

This work was supported by the EU NR 658479 SpikeControl, the Spanish Research Project ER CEREBROT TIN2016-81041-R funded by AEI and FEDER, and the ANR – Essilor SilverSight Chair ANR-14-CHIN-0001.

## Financial interests or conflicts of interest statement

The authors declare that the research was conducted in the absence of any commercial orf inancial relationships that could be construed as a potential conflict of interest

## Author contribution

NRL, ER, and AA conceived the initial idea. FN, RR and NRL designed, modelled and implemented the cerebellar network and the set-up experimentation. NRL and AA prepared figures and drafted the manuscript. All authors reviewed the manuscript and approved the final version.

## Supporting Information

**Figure S3-1.**
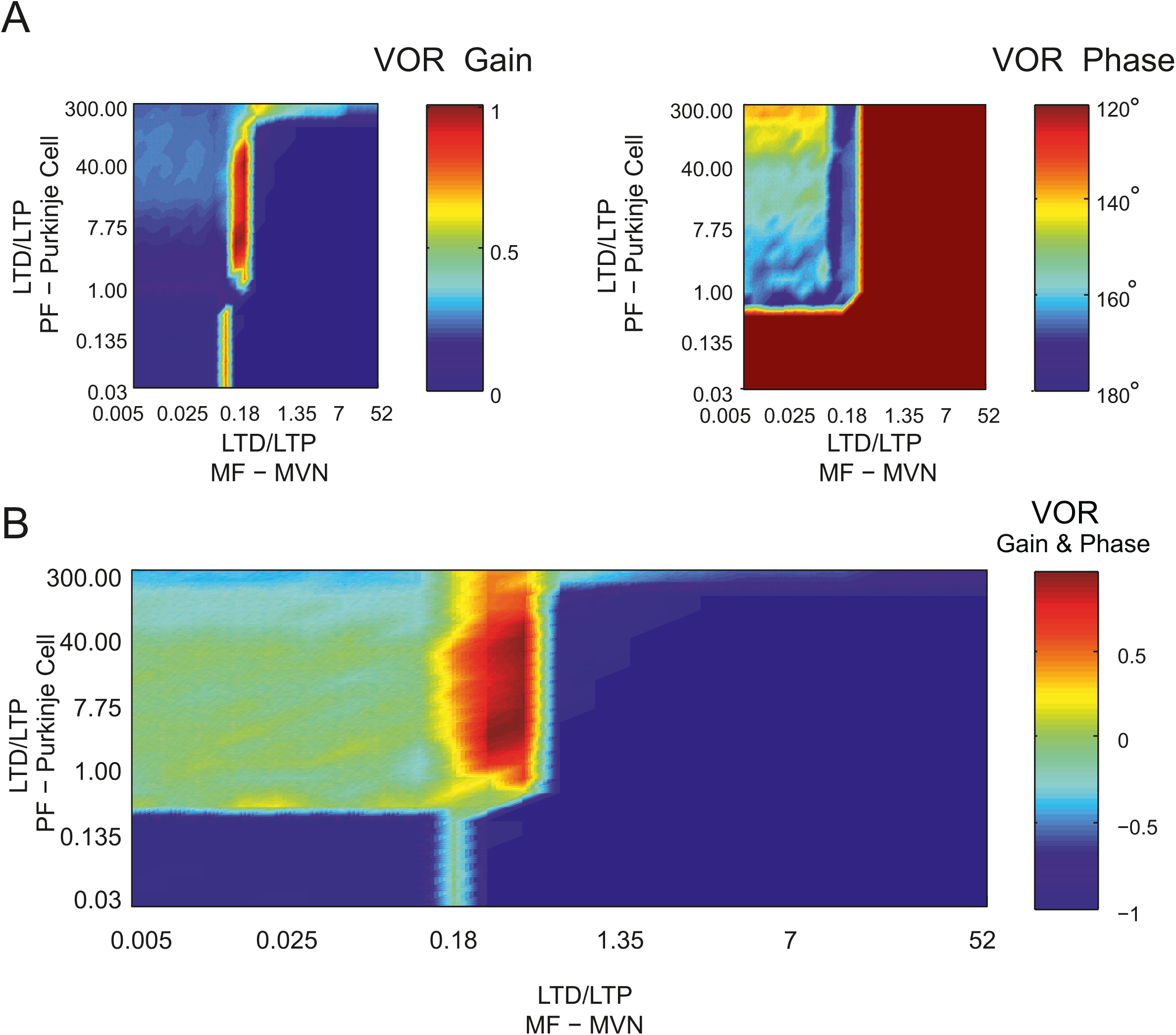
Critical LTD/LTP balance at PF-Purkinje cell and MF-MVN synapses: sensitivity analysis. Cerebellar adaptation modulates PF-Purkinje cell synaptic weights as well as MF-MVN synapses [6, 92]. For synaptic adaptation, the model uses supervised STDP, which exploits the interaction amongst unsupervised and supervised cell inputs to regulate and stabilise postsynaptic activity. Balancing supervised STDP, and the resulting synaptic modification dynamics, is critical, given the high sensitivity of the process that determines the LTD/LTP ratio [127, 128]. A sensitivity analysis of the parameters governing LTD and LTP, shows that LTP exceeding LTD values for a narrow range at MF-MVN synapses preserves VOR learning stability. This holds independently for both VOR gain and phase **(A)** as well as for the combination of the two **(B)**. By contrast, PF-Purkinje cell synapses admit broader limits for the LTD/LTP ratio (A, B). *More detailed description:* we systematically simulated LTP/LTD ratio values at PF-Purkinje cell and MF-MVN synapses within a plausible range that may satisfy the expected h-VOR outcome. As simulations ran, the solutions were iteratively checked until finding the set of LTD/LTP ratio values that exhibited the better performance in terms of h-VOR gain and phase. LTD/LTP balance at each site was modified by systematically multiplying LTD by 1.5^N^ where –11 ≤ N ≤ 12 for PF-Purkinje cell and MF-MVN synapses. For each parameter setting, the cerebellar model underwent 10 000 sec of VOR learning (1Hz head rotation movement to be compensated by contralateral eye movements. **(A)** Final VOR gain and phase plotted over the LTD/LTP range of values that were tested. **(B)** Combined VOR gain and phase (normalised) as a function of the LTD/LTP ratio. At PF-Purkinje cell synapses the LTD/LTP was well balanced for N values ranging between [–1, 7]. At MF-MVN the LTD/LTP balance was more critical since N is within a narrower band range [–1, 0]. The reddish area within the last plot indicates the optimal parameters range. LTP must exceed LTD at MF-MVN synapses for optimal VOR performance. This result is consistent with the unsupervised nature of the LTP for the kernel defined for MF-MVN STDP. Unsupervised LTP with larger values than LTD takes the MF-MVN synaptic weights to the upper bound of their synaptic efficacy, thus provoking more MVN activations. In the absence of LTD counteraction, the cerebellar output is, therefore, upper saturated. LTD driven by Purkinje cell activity blocks LTP at MF-MVN synapses, thus shaping the cerebellar compensatory output.

**Figure S3-2.**
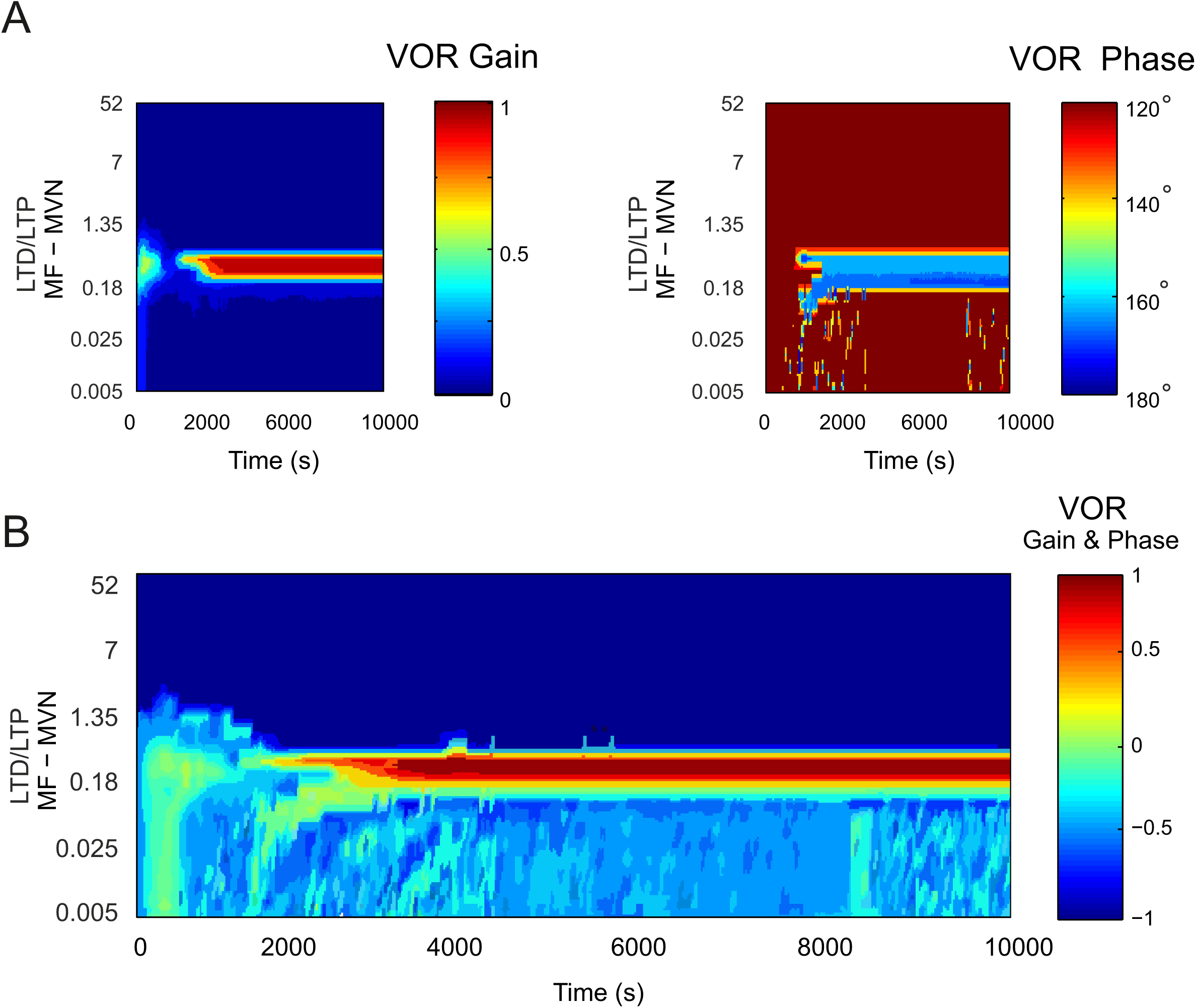
LTD/LTP balance at MF-MVN synapses over time. Whilst LTD/LTP balance was fixed at PF-PC synapses, we modified the LTD/LTP balance at MF-MVN synapses by systematically varying the ratio by 1.5^N^ where –11 ≤ N ≤ 12 during a 10000 sec simulation. **(A)** Final VOR gain and phase plotted as a function of the tested LTD/LTP range across time. **(B)** Combined VOR gain and phase (normalised) over time. A proper balance between LTD and LTP (ratio of approximately 0.4) makes the cerebellum perform optimally after 750 sec.

**Figure S3-3.**
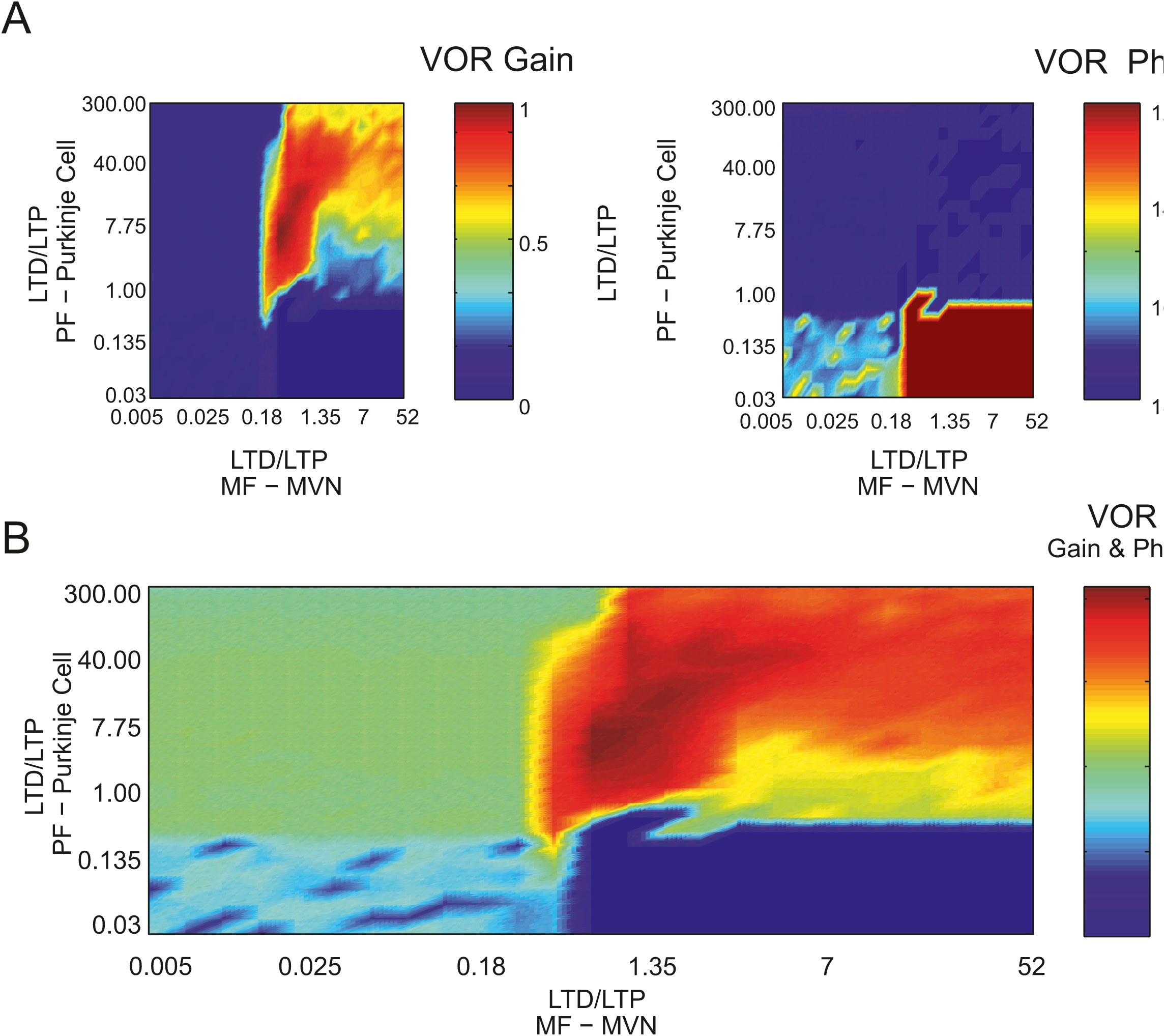
Parameter sensitivity analysis for the LTD/LTP balance at PF - Purkinje cell and MF-MVN synapses in the absence of Purkinje spike burst-pause dynamics. Similar to Fig. 3-1, the parameters regulating the LTD/LTP ratio were exhaustively tested whilst the cerebellar model without Purkinje complex spiking underwent h-VOR learning during a 10000 sec simulation. **(A)** Final VOR gain and phase plotted over the LTD/LTP range of tested values. **(B)** Combined VOR gain and phase (normalised) as a function of the LTD/LTP ratio. LTD/LTP at both PF-Purkinje cell synapses is well balanced for N values ranged between [–1, 7]. Thus, the absence of bursting and pause dynamics leads to a wider range values for the LTD/LTP balance.

**Figure S5-1.**
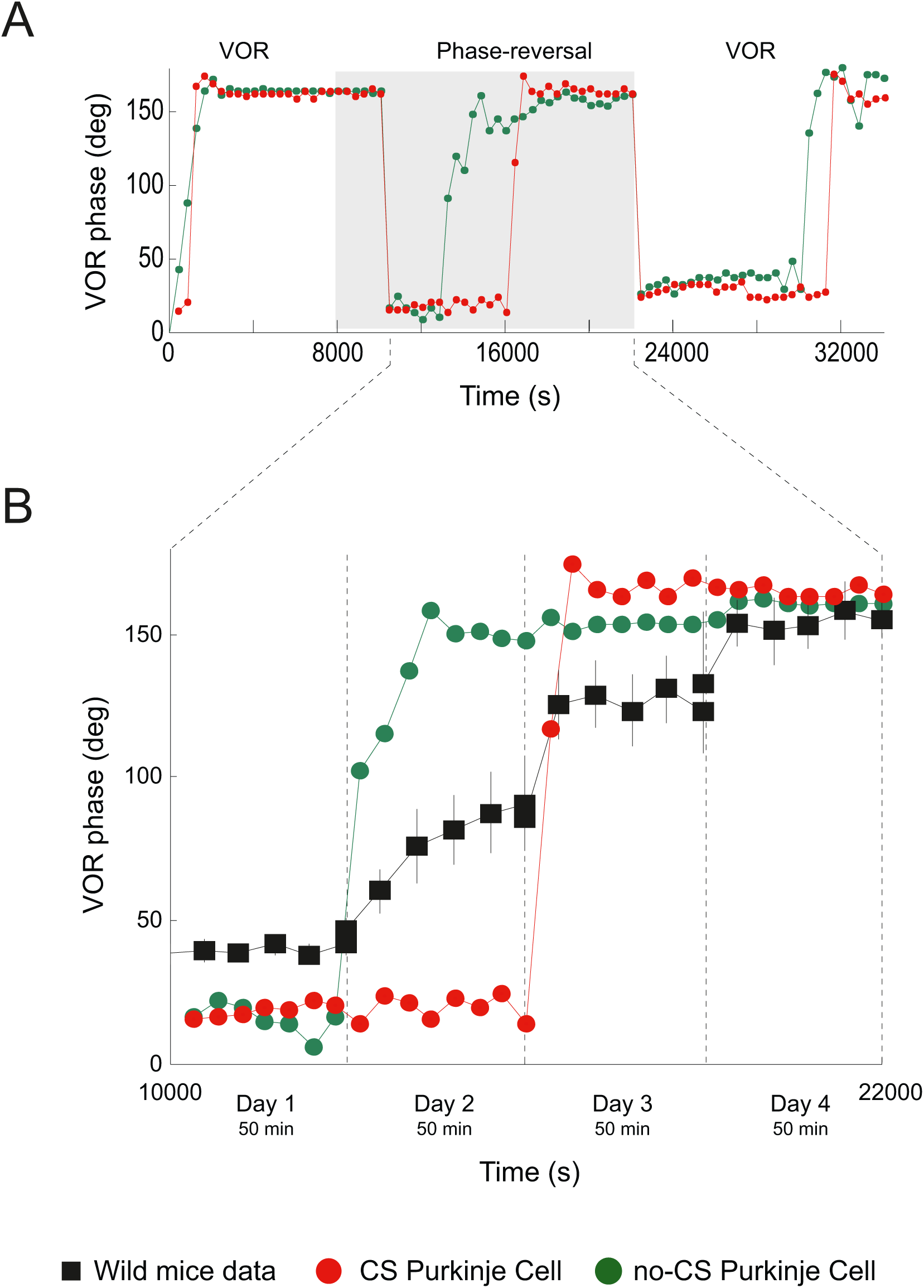
VOR phase-reversal learning: time course of the VOR phase. **(A)** VOR phase adaptation with (red curve) and without (green curve) Purkinje spike burst-pause dynamics. **(B)** Focus is on the phase-reversal period and comparison with experimental data [3].

**Figure S6-1.**
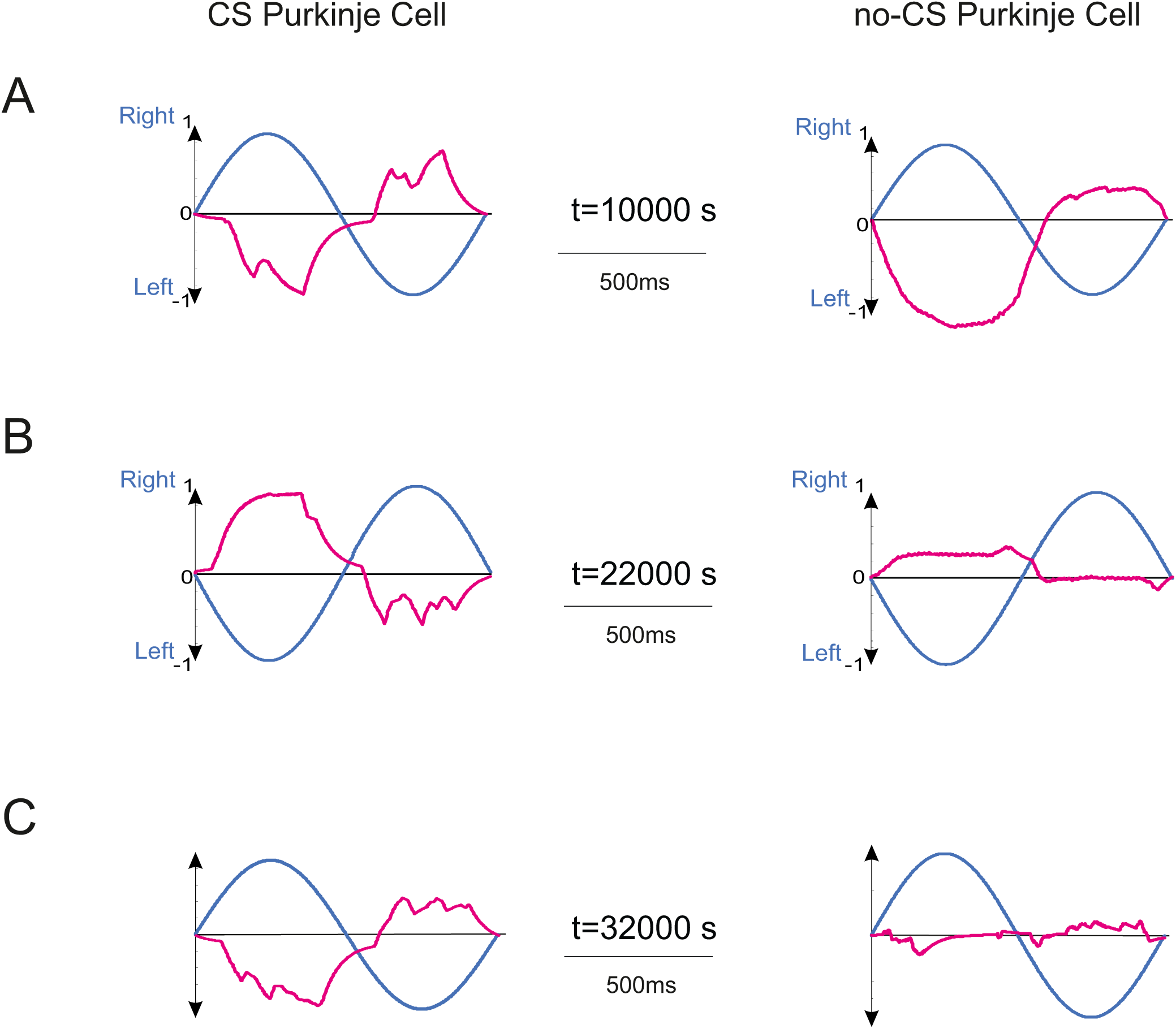
Eye velocity evolution during VOR phase-reversal learning. **(A)** Only the eye velocity movement corresponding to the sparser and more selective distribution of MF-MVN synaptic weights is able to counteract the head velocity movement in counter phase (**B**), as phase-reversal learning is achieved (**C**).

**Figure S7-1.**
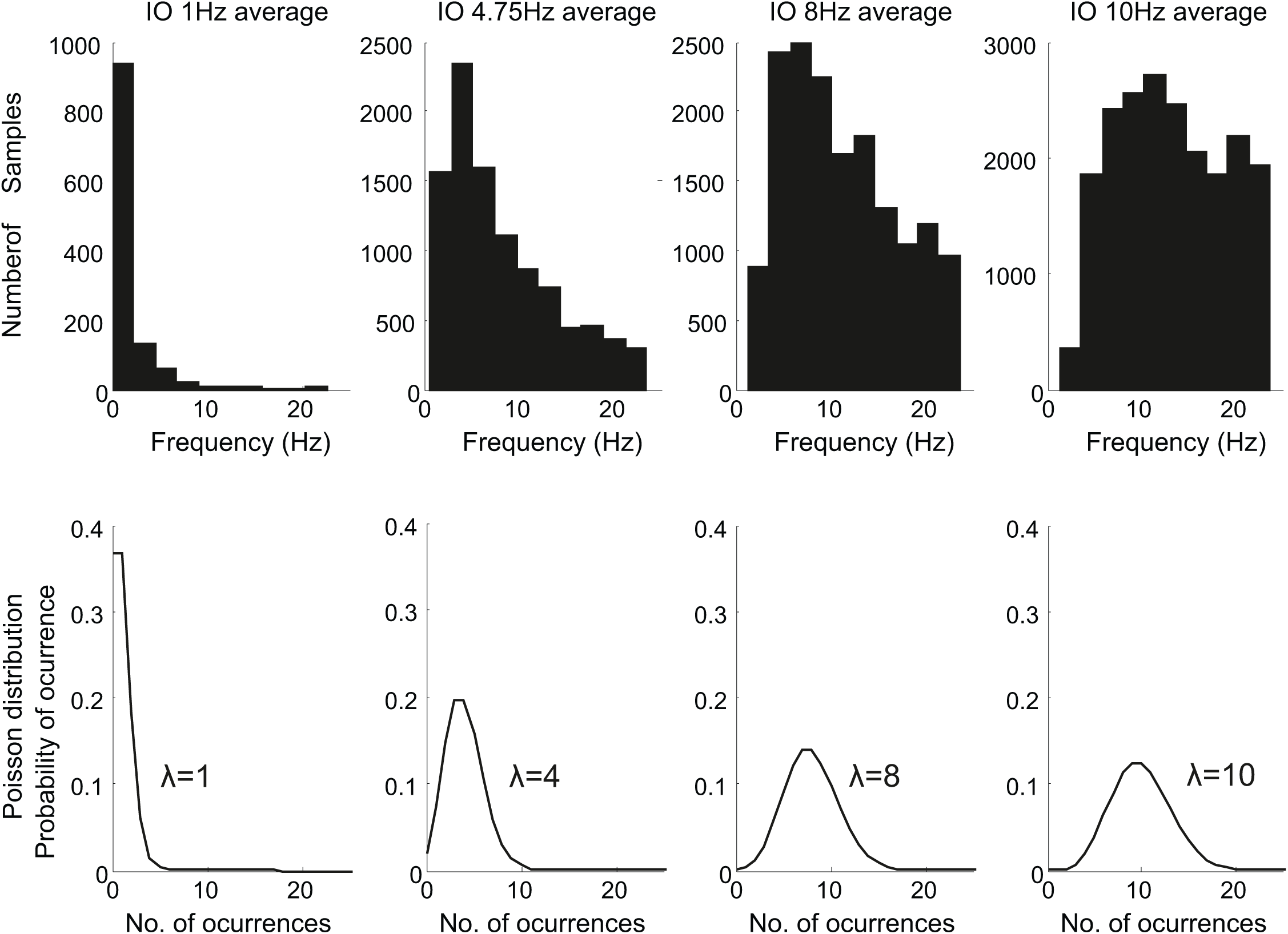
Climbing fibre activation. In the model, CF responses follow a probabilistic Poisson process. Given the normalised error signal ε(t) obtained from the retina slip and a random number η(t) between 0 and 1, the model CF fires a spike if ε(t) > η (t); otherwise, it remains silent[79] A single spike is then able to report timed information regarding the instantaneous error. Furthermore, the probabilistic spike sampling of the error ensures that the entire error region is accurately represented over trials with a constrained CF activity below 10 spikes per second, per fibre (CF activated between 1-10 Hz). Hence, the error evolution is accurately sampled even at a low frequency [115, 117]. This firing behaviour is consistent to those observed in neurophysiological recordings [116].

